# Engineered Lactate Catabolizing Probiotics Reveal Timescale Dependent Microbiome-Host Metabolic Coupling

**DOI:** 10.64898/2026.04.07.716956

**Authors:** Noah T. Hutchinson, Ningyuan Ye, Maria Jennings, Chengyuan Fang, Nathan Qi, Jiahe Li

**Affiliations:** Department of Biomedical Engineering, University of Michigan, Ann Arbor, MI, 48109, United States.; Department of Esophageal and Mediastinum, Harbin Medical University Cancer Hospital, Harbin, Heilongjiang, 150001, China; Department of Molecular and Integrative Physiology, University of Michigan, Ann Arbor, MI, 48109, United States

## Abstract

The exchange of lactate, a metabolic substrate and regulator, between the gut lumen and systemic circulation for use in host and microbial processes is well documented, but tools capable of uncovering whether this process influences host metabolic status across acute and chronic contexts are lacking. In our prior work, we engineered probiotic *Bacillus subtilis PY79* to produce lactate oxidase (LOX) intracellularly, allowing it to rapidly convert intestinal lactate to pyruvate. Following oral administration, LOX reduced systemic lactate concentrations at rest and under challenge conditions, providing a platform for investigating lactate’s influence on host metabolism and microbiota. In the present work, we demonstrate that acute LOX administration effectively rewired microbiota function and host energy balance, as revealed by 16S sequencing and indirect calorimetry. *In silico* microbial community modeling via MICOM and metagenomic inference via PICRUSt2 suggested that acute shunting of lactate to pyruvate induced microbiota remodeling towards anabolic processes, reflected by increased flux of pyruvate, acetate, and formate, alongside moderate to large increases (Cohen’s d = 0.60-1.00) in pathways for fructan degradation, B-vitamin biosynthesis, and lipid synthesis. These anabolic shifts temporally aligned with transient increases in host energy expenditure (β = 1.08, p<0.05) via glucose oxidation (β = 0.01, p<0.05), hinting at functional coupling between microbial biosynthesis and host energy balance via lactate exchange. Of note, acute LOX administration also improved thermoregulation and survival following LPS-induced sepsis, demonstrating functional relevance of these metabolic effects during acute inflammatory challenge. To assess chronic effects, we administered LOX for 6 weeks during diet-induced obesity. LOX treatment persistently reduced blood lactate. However, this chronic lactate reduction did not curtail the progression of diet-induced obesity or induce sustained modulation of host energy expenditure. This disconnect between acute and chronic findings suggests that gut-centric lactate conversion affects energy balance through microbiome and/or host-dependent mechanisms, but cannot override homeostatic forces in the long term to produce clinical benefit during chronic disease. Our results validate LOX probiotics as a tool for acute metabolic augmentation, and highlight a clear homeostatic limit to gut-centric therapies. This platform may enable targeted design of probiotic interventions matched to therapeutic timescale and inform synbiotic formulations that overcome homeostatic compensation.

## Introduction

The gut microbiota functions as a metabolic interface between the host and its environment, contributing to energy harvest from food, production of gut hormones, metabolic signaling, and immune programming (1, 2). Consequently, the structure and function of the gut microbiota has been linked to the progression of acute and chronic diseases, with preclinical studies drawing causal connections (3–6). Microbial metabolites such as SCFA contribute to whole body energy expenditure, and are associated with critical illness related outcomes (7–9), and dysbiosis of these metabolic networks is implicated in conditions ranging from metabolic disease to acute illnesses such as sepsis (5, 10–18). Conversely, targeted microbiome therapies can either prevent these conditions or improve outcomes (19–23). Despite these associations, tools for targeted manipulation of specific metabolic nodes within this host-microbiome axis remain lacking, limiting mechanistic understanding and rational therapeutic design.

Given its broad network of metabolic influence and functional adaptability, a growing body of research has focused on leveraging the gut microbiome to elicit functions that treat metabolic dysfunction, using probiotics, prebiotics, postbiotics, symbiotics, and fecal microbiota transplantation (24–37). More recently, preclinical studies have demonstrated that genetically engineered bacteria can modulate host metabolism in rodent models by synthesizing exogenous molecules, including GLP-1 receptor agonists (38–40), anorexigenic lipids (41), and fermentative end products such as butyrate (42). Together, these natural and engineered live biotherapeutics illustrate the potential of gut-resident bacteria as factories for in situ delivery of peptides, proteins, and other biologics. However, the sustained synthesis of potent host signaling molecules presents substantial safety and regulatory challenges related to uncontrolled dosing, biocontainment, and side-effect profiles.

An alternative strategy is to target metabolites that lie at the interface of host and microbial metabolism. Recent evidence in elite athletes provides a compelling example. It was found that fecal sampling of Boston marathon runners around race time revealed increased colonization by lactate-catabolizing microbes and an enriched capacity to utilize lactate as a carbon source to generate beneficial metabolites (43). Given the known associations between lactate accumulation, cardiovascular fitness, metabolic dysfunction, and acute illness (44–51), this bidirectional relationship suggests that lactate may be a tractable metabolic node for microbiota induced modulation of host energetic status. With that said, the contribution of local lactate conversion to microbiota bioenergetic output and downstream host physiology remains poorly understood due to the lack of viable tools for targeted manipulation of microbial lactate utilization. While reduction of the systemic lactate: pyruvate ratio via enzyme administration has been shown to improve tissue-level redox status (52), an approach that provides native gut bacteria direct access to the resultant pyruvate has yet to be explored. Our prior studies, in which we engineered *Bacillus subtilis* PY79 to express lactate oxidase and rapidly convert lactate to pyruvate, demonstrate that oral administration of this strain reduces systemic lactate levels both at rest and under challenge conditions (53). With this refined model, we can directly interrogate how targeted lactate-to-pyruvate conversion in the gut influences both microbiota metabolic function and host physiology.

Using our established model, which utilizes the gut and its microbiota as a metabolic “factory” for local and systemic lactate conversion via an engineered probiotic, we characterized its effects on the microbiome’s metabolic profile and host metabolism across acute and chronic timescales. We combined next-generation sequencing, bioinformatic tools, and indirect calorimetry to assess acute effects on microbiome remodeling and host energetics, tested whether these acute effects confer protection during LPS induced sepsis, and evaluated whether chronic administration could sustain metabolic benefits during diet induced obesity. This functionally driven strategy enables the precise interrogation of the role of lactate in microbiome–host metabolic coupling, characterizes the temporal dynamics of single-point metabolite interventions, and sets a rational foundation for mechanistically targeted multi-component strategies.

## Materials and Methods

### Chemicals and Reagents

All chemicals and bacterial culture broth were purchased from Fisher Scientific International Inc. (Cambridge, MA, USA) unless otherwise noted and were of the highest purity or analytical grade commercially available. All molecular cloning reagents, including restriction enzymes, competent cells, and the HIFI assembly kit, were purchased from New England Biolabs (Ipswich, MA, USA). DNA oligonucleotides were ordered from Sigma-Aldrich (St. Louis, MO, USA). gBlocks were ordered from IDT with *B. subtilis* codon optimization.

### General Growth Conditions for B. subtilis

*B. subtilis* strains were grown at 30 °C and 220 rpm overnight in LB medium with or without 5 μg/mL chloramphenicol. For downstream applications, overnight bacterial culture was diluted 10 times, followed by culturing at 37 °C and 220 rpm until the OD_600_ reached between 0.6 and 0.8.

### Strain Construction

Strain construction is described in our previous manuscript (53). All *B. subtilis* strains were constructed in PY79. All constructs were confirmed by Sanger sequencing.

All *amyE* locus integration fragments contain three parts: *amyE* 5′ arm with *P_NBP3510_*, GOI, and *amyE* 3′ arm with chloramphenicol resistance. The integration fragment was assembled from different plasmids and chromosomal DNA of *B. subtilis* to make *ΔamyE::cat*_sfGFP first, which was used as a new template to later make LOX constructs. A FLAG tag was added at 3′ of LOX.

### ΔamyE::cat_sfGFP

The *amyE* 5′ arm was amplified through WB800N genomic DNA with the primers B685_V2/B742. *P_NBP3510_*_sfGFP was amplified from WBSGFP with B741/B747. *amyE* 3′ arm was amplified through lab plasmid pHT01-NBP3510-sfGFP with B746/B690. Three fragments were first combined through a HIFI assembly kit followed by cementing PCR before transformation into *B. subtilis* as described below.

### ΔamyE::cat_LOX

*amyE* 5′ arm with *P_NBP3510_* and *amyE* 3′ arm with Cm^r^ were amplified separately through the chromosomal DNA of PY79 *ΔamyE::cat*_sfGFP.

### Spore Preparation

Overnight bacterial culture was plated on selective 2×Schaeffer’s medium-glucose plates and incubated at 30 °C for 72h (54). Spores were scraped from the plate into centrifuge tubes and then sonicated with a metal probe at 20 W for three cycles of 1 minute with a 1-minute rest in between. Following sonication, spores were pelleted via centrifugation at 14,000 rpm for 30 min at 4 °C and then washed and resuspended in ddH_2_O. This process was repeated once daily for three days. Spores were aliquoted at 2×10^10^ CFU/mL, stored at −20 °C, and thawed upon use.

### General Conditions and Practices for Mouse Experiments

All animal experiments were approved by the University of Michigan Institutional Animal Care and Use Committee (IACUC) and were conducted in AALAC-accredited facilities. C57BL/6J mice were purchased from Jackson Laboratories and were all acclimated for 1-2 weeks prior to experimental use. During acclimation, the dirty bedding from all cages was combined, mixed, and evenly distributed among all cages to homogenize the baseline microbiome composition.

### Genomic DNA Extraction from Feces

C57BL/6 mouse feces were collected before the first gavage and 1, 3, and 7 days after the first gavage. Feces were immediately frozen in liquid nitrogen and saved in −80 °C for later use. Genomic DNA was extracted using the Quick-DNA Fecal/Soil Microbe Miniprep Kit (Zymo, catalog # D6010) according to the manufacturer’s instructions.

### 16S rRNA Sequencing

The V4 region of the 16S rRNA gene was amplified with specific primers (5′-GTGCCAGCMGCCGCGGTAA-3′ and 5′-GGACTACHVGGGTWTCTAAT-3′). All PCR reactions were carried out in 30 μL reactions with 15 μL of Phusion High-Fidelity PCR Master Mix (New England Biolabs), 0.2 μM forward and reverse primers, and about 10 ng of template DNA. Thermal cycling is initiated with an initial denaturation step at 98 °C for 1 min, followed by 30 cycles of denaturation at 98 °C for 10 s, annealing at 50 °C for 30 s, and elongation at 72 °C for 60 s. Finally, the solution was kept at 72 °C for 5 min. PCR products were mixed with an equal volume of SYBR Green loading buffer and subjected to electrophoresis on 2% agarose gel for quantification and qualification, followed by purification with the GeneJET Gel Extraction Kit (Thermo Scientific). Sequencing libraries were generated using the NEB Next Ultra DNA Library Prep Kit for Illumina (NEB, USA) according to the manufacturer’s recommendations, and index codes were added. The library quality was assessed on a Qubit@ 2.0 fluorometer (Thermo Scientific) and an Agilent Bioanalyzer 2100 system. At last, the library was sequenced on an Illumina HiSeq 2500 platform, generating 250 bp paired-end reads.

### Microbiome Data Analysis

Dada2 preprocessing (55) was used on raw sequence files generated by Novogene. Based on qualitative assessment of quality plots, forward reads were trimmed at 240 bp and reverse reads were trimmed at 160 bp to ensure uniformity and preserve read quality. After dereplication, denoising, merging, and chimera removal, the Silva 138.1 database was used to assign taxonomy (56, 57). Following merging with metadata and a random phylogenetic tree, α diversity was calculated via Faith’s PD (phylogenetic diversity), Chao1 index (community richness), and Shannon index (more sensitive to evenness). Downstream from this, Amplicon sequence variants (ASVs) not present in two or more samples at a frequency of 11% or more were deemed to be noise and removed from the analyses. To understand the nature of compositional shifts, β-diversity distance matrices were calculated using weighted UniFrac, Bray-Curtis, and Aitchison distances. A centered log-ratio transformation was applied prior to calculating Aitchison distances to account for compositionality. Principal coordinate and principal component analyses were conducted and plotted for weighted UniFrac/Bray-Curtis and Aitchison distances, respectively. For statistical comparisons of α and β diversity, a linear mixed-effects modeling approach was employed in a dose- and time-aware manner, which included binary fixed effects for both probiotic administration and the inclusion of the LOX enzyme, as well as consideration of time relative to treatment, cumulative doses of both LOX and probiotic, and a random effect for subject ID (see **Table 1**). For differential abundance testing, we used this model along with a complementary discovery model with binary effects for LOX and probiotic to maximize statistical power and identify broader groups of responsive taxa. As a result, the dose-aware model could be used to distinguish binary treatment effects from dose-response relationships. MaAsLin2 was used to calculate differentially abundant ASVs (58), adonis PERMANOVA testing was used to compare β diversity (59, 60), and a regular LMM was applied to compare α diversity metrics.

**Table 1:** Model Variables by Treatment Group for Bioinformatic Analyses.

| Treatment Group | Baseline | Early | Mid | Late | pro | lox |
| --- | --- | --- | --- | --- | --- | --- |
| PBS | n=4 | n=4 | n=5 | n=5 | absent | absent |
| GFP (Probiotic) | n=5 | n=5 | n=4 | n=5 | present | absent |
| Lox (Probiotic + Lox) | n=5 | n=5 | n=5 | n=5 | present | present |

***Cumulative Dose Values by Timepoint***
| Treatment Group | Baseline (0h) | Early (24h) | Mid (72h) | Late (D7) |
| --- | --- | --- | --- | --- |
| PBS - cumulative_doses_pro | 0 | 0 | 0 | 0 |
| PBS - cumulative_doses_lox | 0 | 0 | 0 | 0 |
| GFP - cumulative_doses_pro | 0 | 1 | 2 | 3 |
| GFP - cumulative_doses_lox | 0 | 0 | 0 | 0 |
| Lox - cumulative_doses_pro | 0 | 1 | 2 | 3 |
| Lox - cumulative_doses_lox | 0 | 1 | 2 | 3 |

Total n = 57 samples ***Model Term Definitions and Interpretations***
| Model Term | What It Compares | Which Groups | Biological Interpretation |
| --- | --- | --- | --- |
| pro | Effect of probiotic presence | GFP+Lox vs PBS | Does having probiotic bacteria alter community structure? |
| lox | Effect of lox presence | Lox vs GFP+PBS | Does having lox bacteria alter community structure? |
| cumulative_doses_pro | Linear dose effect of probiotic | Increasing doses within GFP+Lox groups (0>1>2>3) | Does more probiotic exposure lead to greater community changes? |
| cumulative_doses_lox | Linear dose effect of lox | Increasing doses within Lox group (0>1>2>3) | Does more lox exposure lead to greater community changes? |
| treatment_phase | Time effects | All timepoints compared to baseline | Does the microbiome change over the experimental timeline independent of treatments? |

### 16S metagenome prediction and downstream analyses

Processed sequences were used for metagenome prediction and downstream analyses via the widely used Phylogenetic Investigation of Communities by Reconstruction of Unobserved States 2 (PICRUSt2) software (61). Since this was done on a Windows device, WSL/Ubuntu was used to open a virtual RStudio in a Linux interface, where an R Markdown file was used to execute all functions, which infers the calculation of predicted metagenomes. Once output files were generated, they were exported for downstream analyses via desktop Rstudio, with statistical modeling being the same as was used for microbiome analyses (**Table 1**). Predicted pathway abundances were used to minimize noise in the data. Pathways with fewer than 10 total reads, an average relative abundance of less than 1 × 10^5^, and not present in at least 10% of the samples were considered noise and were filtered out. Filtered data were all CLR transformed prior to downstream analyses, with a pseudo count of 0.5 to avoid issues with 0 values. Aitchison distances were calculated for PCA and comparison via PERMANOVA, and MaAsLin2 was used to determine global differential abundance. The top 10 contributing features to PC1 and PC2 were identified, and these features were further analyzed using LMM with time, treatment, and the time × treatment interaction as fixed effects. Given the targeted and exploratory nature of this analysis, p-value corrections were not used for the sake of generating hypotheses.

### Gut Microbiome Metabolic Modeling

Community-level metabolic modeling was performed using MICOM (v0.37.1) (62) with the AGORA2 genus-level model database (agora201_refseq216_genus_1) (63). To maximize AGORA2 mapping coverage, taxonomy was reassigned from the aforementioned DADA2 outputs using Greengenes2 and processed to genus-level relative abundance (ref, mean 96% of reads mapped per sample). All three treatment groups were analyzed (PBS, sfGFP, LOX, n=18-20 samples per group across all timepoints: 0h, 24h, 72h, D7). Two simulation conditions were modeled: 1) native simulations using the observed community composition from 16s sequencing for all 57 samples, 2) simulations in which *B. subtilis* was added at 3% relative abundance, either with a forced L-lactate uptake flux of −12.0 mmol/gDW/h to model LOX enzymatic activity (for LOX samples), or without LOX function (for sfGFP samples). In the latter, PBS simulations were still run on the native community. Due to the lack of evidence regarding the specific metabolic pathways engaged within *B. subtilis* PY79 engineered to express lactate oxidase, L-lactate uptake was constrained as a proxy for LOX enzymatic activity, and downstream metabolic routing within B. subtilis was determined by the AGORA2 metabolic model. An EU average diet was applied as a proxy for standard mouse chow, as no mouse-specific dietary constraint was available in this format. Cooperative tradeoff optimization was set to a tradeoff value of 0.9. Net community exchange fluxes for key metabolites (pyruvate, acetate, formate, butyrate, succinate, L-lactate) were extracted and compared between groups. Between-group comparisons at 72h were performed using unpaired t-tests, and within-animal enzymatic effects (native vs. with LOX) were assessed using paired t-tests. All comparisons are exploratory given the sample size (n = 5/group per timepoint), so no correction for multiple comparisons was applied.

### Treadmill Running, Concurrent Indirect Calorimetry, and Blood Lactate Measurements

The study design for treadmill experiments is displayed in **Figure 4A**. Briefly, once daily doses of 2×10^9^ CFU were administered to female C57BL/6J mice at 9am for three days. Twenty minutes after the third dose, a baseline blood lactate measurement was taken, and mice were placed in their respective treadmills. Twenty mice were divided between a traditional treadmill, where blood lactate measurements could be taken immediately following exercise, and a treadmill chamber integrated into an open-circuit calorimeter, where mice were kept inside the chamber for an additional 5 minutes after running to characterize their metabolic status during recovery. Furthermore, each group of 10 mice was divided into either the sfGFP or LOX group (n = 5/group). In mice undergoing indirect calorimetry, blood lactate measurements were taken 5 minutes after exercise cessation. In both groups, an additional blood lactate measurement was taken 20 minutes after exercise ceased.

Oxygen consumption (VO2) and carbon dioxide production (VCO2) were measured using the Comprehensive Laboratory Animal Monitoring System (CLAMS, Columbus Instruments), an integrated open-circuit calorimeter, consistent with prior studies of indirect calorimetry in rodents [1,4,6]. Before the study, the mice were each placed into the treadmill chambers to acclimate them to the treadmill environment for 10 minutes. For 2 days prior to the study, the mice were individually put on the same treadmill for 30 minutes each day. Mice were weighed prior to the running test. They were then individually placed into the sealed treadmill chambers (305 x 51 x 44 mm³). The slope of the treadmill was set at 15° to the horizontal throughout the study. The study was conducted in an experimental room maintained at 20–23 °C with 12-hour dark-light cycles (6:00 PM–6:00 AM). The measurements were only carried out between 9:00 AM and 2:30 PM on each day. During this time, the animals were run on the treadmills one at a time, and the treadmill was wiped clean between each test. The system was routinely calibrated before the experiment using a standard gas (20.6% O2 and 0.5% CO2 in N2). VO2 and VCO2 in each chamber were sampled continuously at 5-second intervals. The airflow rate through the chambers was set at 0.80 L/min. Respiratory exchange ratio (RER) was calculated as VCO2 / VO2, following established equations for substrate oxidation (64, 65). Total energy expenditure, carbohydrate oxidation, and fatty acid oxidation are calculated, respectively, based on the values of VO2 and VCO2 using equations as listed in the data sheets. The protein breakdown (which is usually estimated from urinary nitrogen excretion) was ignored (66).

All the mice were ran under the same standard treadmill schedule, which was:

- 10 minutes baseline recording
- 5 minutes @ 5 m/min
- 5 minutes @ 9 m/min
- 5 minutes @ 12 m/min
- 5 minutes @ 15 m/min
- 2 minutes @ 17 m/min
- 2 minutes @ 19 m/min
- 2 minutes @ 21 m/min
- 2 minutes @ 23 m/min
- 2 minutes @ 25 m/min
- 2 minutes @ 27 m/min
- 2 minutes @ 29 m/min
- 2 minutes @ 31 m/min
- 2 minutes @ 33 m/min
- 2 minutes @ 35 m/min
- 2 minutes @ 37 m/min
- 2 minutes @ 39 m/min
- 2 minutes @ 41 m/min
- 2 minutes @ 43 m/min
- 2 minutes @ 45 m/min
- 2 minutes @ 47 m/min

Exhaustion was qualified as a mouse sitting on the shocker (1.60mA, 120v, 3Hz) for 5 consecutive seconds, at which point the shocker was shut off, the treadmill schedule stopped, and for mice undergoing indirect calorimetry measures, they were kept in the chamber for an additional 5 minutes to generate recovery data before the animal was removed from the treadmill. Use of treadmill testing for murine exercise capacity and endurance phenotyping is well-established (67–69).

### Metabolic Cage Assessments

In experiments conducted on young, healthy mice, a similar study design was employed, utilizing a treadmill setup with the addition of blood lactate measurements. Mice were given once-daily doses of 2 × 10^9^ CFU at 9am for three days, and then placed in the metabolic cage system 20 minutes after the third dose. For the experiment with obese mice, where they were dosed once every three days, 5 weeks into the experiment, mice were given a dose in the morning and placed in the cages an hour later. Mice were kept in the cages for three days.

Oxygen consumption (VO₂), carbon dioxide production (VCO₂), and spontaneous motor activity were measured using the Promethion (Comprehensive, High-Resolution Behavioral Analysis Systems, Sable Systems International), an integrated open-circuit calorimetry system equipped with an optical beam activity monitoring device. Mice were weighed each time before the measurements and individually placed into the Mouse Cage (Model 3721; 8.1 × 14.4 × 5.5 in.) with free access to food and water. The study was conducted in an experimental room maintained at 20–23 °C with 12–hour (6:00 PM–6:00 AM) dark-light cycles. The measurements were carried out continuously for 120 hours. During this time, animals were provided with food and water through the equipped feeding and drinking devices located inside the cage. The Promethion food, water intake, and body weight monitoring system features high-precision sensors capable of measuring real-time data for mice and rats. The system was routinely calibrated each time before the experiment using a standard gas (20.5% O₂ and 0.5% CO₂). VO₂ and VCO₂ in each cage were sampled sequentially for 30 seconds in a 5-minute interval, and motor activity was recorded every second in X, Y, and Z dimensions.

The respiratory quotient (RQ), also known as the respiratory exchange ratio (RER), was calculated as VCO₂/VO₂ (64, 70). Total energy expenditure, carbohydrate oxidation, and fatty acid oxidation can be calculated, respectively, based on the values of VO₂, VCO₂, and the protein breakdown (which is usually estimated from urinary nitrogen excretion) (65–67).

### Diet-Induced Obesity Studies

For obesity modeling, male C57BL/6J mice were purchased from JAX and fed a diet composed of 60% kcal fat, and 20% kcal of both carbohydrate and protein (D12492, Research Diets) ad libitum. Water access was also ad libitum, and mice were housed in thermoneutral conditions (30C).

After two weeks of acclimation to their facility, mice were randomized by body weight to either water (vehicle control), sfGFP (probiotic negative control, 10^11^ CFU/kg), LOX (treatment group, 10^11^ CFU/kg), or semaglutide (positive control and industry benchmark, 40 µg/kg, Medchemexpress cat# HY-114118) treatment. Each treatment was administered once every 3 days. Oral treatments (water, sfGFP, LOX) were administered in 150-250 µL gavages to match the required dose in a 2 × 10^10^ CFU/mL suspension. Semaglutide was prepared and dissolved according to the manufacturer’s instructions and administered via 150 µL IP injections.

### Biodistribution and Fecal CFU Quantification

For biodistribution analyses, mice were euthanized via CO2 asphyxiation and subsequent cervical dislocation. The intestinal regions were removed and separated, and their contents were scraped and added to 300 μL of sterile PBS. For fecal CFU, fresh fecal samples were collected and placed in 300 μL of sterile PBS, followed by a 15-minute incubation. In both cases, the mixture was homogenized using a vortex and then applied to LB agar supplemented with 5 μg/mL chloramphenicol. Plates were incubated at 30°C for 12–16 hours. CFU/g fecal matter was determined by a standard CFU counting method.

### Statistical Analyses

For *in vivo* experiments, repeated-measures data over time were analyzed using linear mixed-effects models. For calorimetry data, mixed-effects models included time (in hours or days) and treatment as fixed effects, along with relevant covariates (e.g., body weight, lean body mass, and blood lactate concentration, when available), and a random intercept for subject. Area under the curve (AUC) was calculated using the trapezoidal method and analyzed via Student’s t-test or one-way ANOVA, as appropriate. AUC values were calculated as aggregate values in GraphPad Prism (v.10.4.1) but plotted as individual values for visualization. In some cases, pairwise comparisons were performed using a t-test for unpaired data in GraphPad Prism (v.10.4.1). Otherwise, all statistics were conducted in RStudio (v.4.3.3), using the lme4 and lmerTest packages. Assumptions of normality and homoscedasticity were assessed prior to model selection. Results from the exploratory temporal trajectory analysis are reported as unadjusted p-values and should be interpreted as hypothesis-generating.

### LPS Induced Sepsis Experiments

Obese male C57Bl/6J mice from JAX (strain #380050) fed the aforementioned high fat diet for 12 weeks with ad libitum access to water were utilized for this experiment. Obese mice were used because obesity is associated with elevated circulating lactate due to increased adipose tissue lactate production (46, 48), providing a physiological context in which LOX-mediated lactate reduction has greater substrate availability and potential therapeutic relevance. Following the same pretreatment process as other acute experiments, mice were administered 5 mg/kg LPS from *Escherichia coli* O111:B4 (Sigma L2630). To account for probiotic excretion via septic diarrhea, additional doses were administered 8h and 24h post LPS. At 24h, blood lactate and body temperature were measured, and survival was monitored every 12h for 5 days. Diet gels and water bottles were provided uniformly upon dosing of LPS.

## Results

### Acute LOX administration drives temporal remodeling of gut microbiota composition and predicted function

To evaluate whether the acute conversion of lactate to pyruvate concurrently induces measurable remodeling of gut microbiota composition and predicts function, we employed comprehensive analyses of 16S rRNA sequencing data from our previously established acute dosing paradigm (53), utilizing expanded bioinformatic approaches, including predicted metagenomic profiling. To distinguish treatment effects from temporal changes and assess whether effects were driven by treatment presence versus cumulative doses, we applied both a binary exposures model and a dose-aware model to parse out effects due to overall treatment, dose-response effects, and time-dependent changes. Despite biodistribution analyses indicating significant CFU detection of both sfGFP and LOX bacteria in the cecum and colon (**Fig. S1**), the effects on microbial composition and predicted functions were subtle, with no observable shifts at the phylum level (**Fig. 1B**). Beta-diversity analyses of Bray-Curtis dissimilarity showed that community compositions changed over time, and both probiotic and LOX administration contributed independently and additively to variance in community structure. In the binary exposures model, both probiotic exposure (R^2^ = 0.061) and LOX inclusion (R^2^ = 0.063) independently altered community structure. Importantly, cumulative LOX doses showed a dose-dependent effect (R2=0.026), suggesting progressive community remodeling with repeated exposure (**Fig. 1D**). These findings were corroborated by weighted UniFrac analyses, which showed similar patterns (probiotic: R²=0.057, p=0.001; LOX: R²=0.077). Alpha diversity, calculated as Shannon Index (evenness), Chao1 Index (richness), and Faith’s PD (phylogenetic diversity), was not affected by timepoint of collection or treatment (**Fig. 1C**). Binary modeling in differential abundance analysis showed the compositional effects of probiotic and LOX to be largely concentrated within the *Firmicutes* phylum. LOX administration altered 25 ASVs within the *Lachnospiraceae* family (11 increased, 14 decreased), whereas probiotic (sfGFP) administration had a smaller effect, with only 9 ASVs altered (3 increased, 6 decreased) (**Fig. 1E, S2**). Dose aware modeling revealed that cumulative doses of LOX led to increases in 3 ASVs considered to be beneficial ‘next generation probiotics’ (2 *Roseburia* and one *Oscillospiraceae*), with cumulative doses of probiotic further contributing to increased *Oscillospiraceae* and *Clostridium* but resulting in decreased *Ligilactobacillus.* Together, these findings suggest that the acute lactate-to-pyruvate conversion results in modest, family-level restructuring of the gut microbiota, without significant changes in overall diversity, with some effects accumulating progressively due to repeated treatment exposure.

**Fig. 1.**
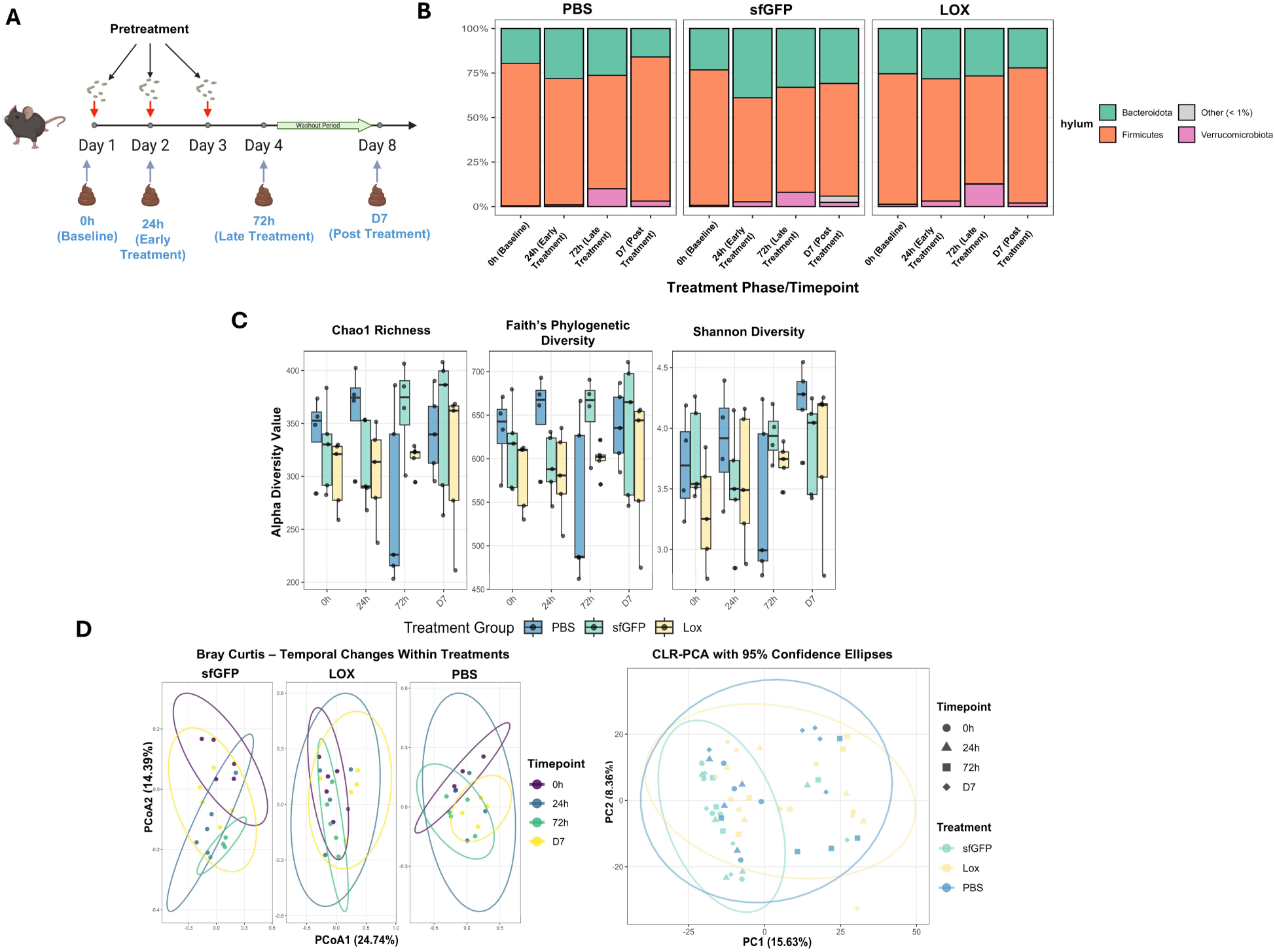
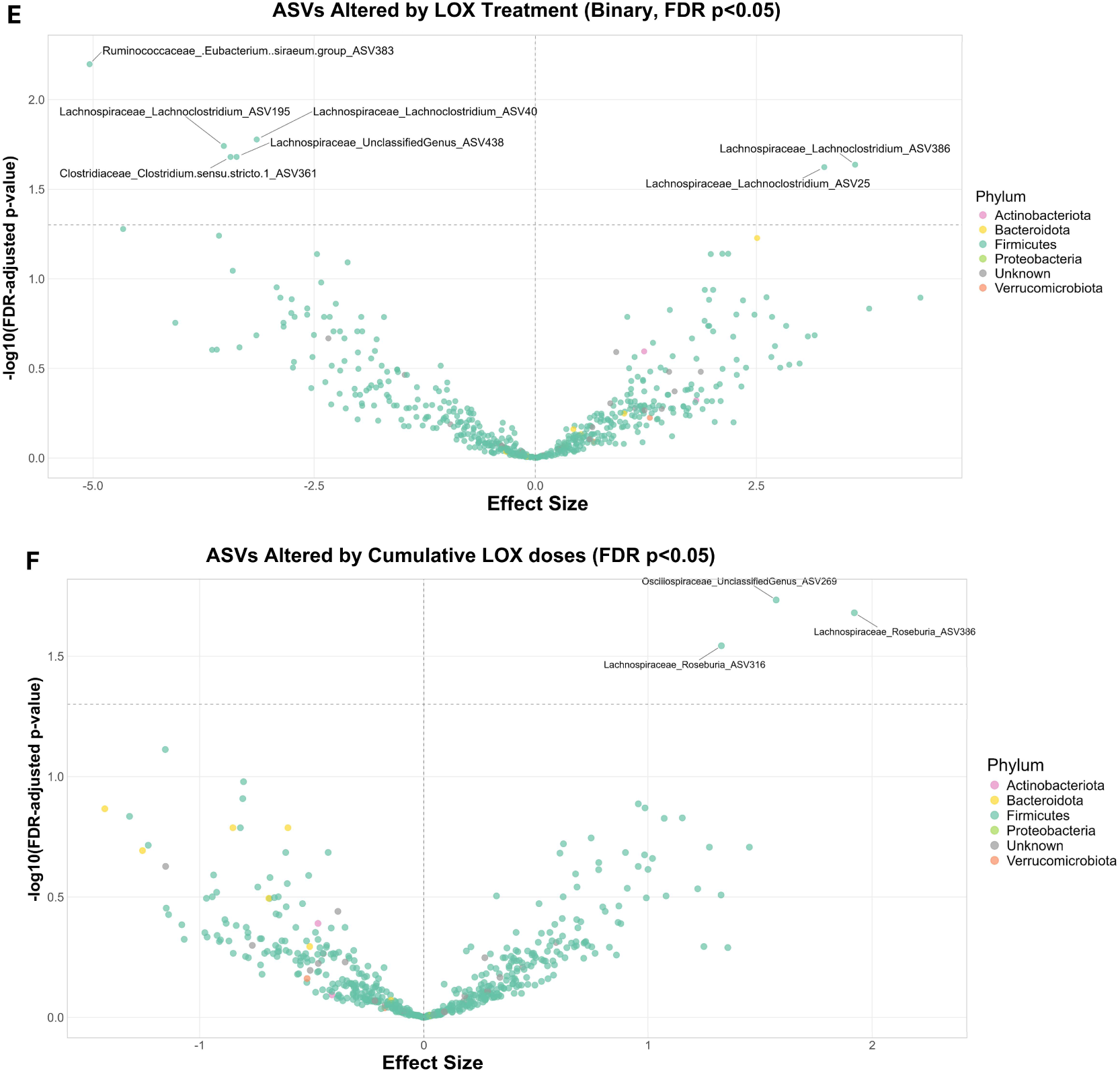
Acute LOX administration drives temporal remodeling of microbiota community composition. (A) Schematic for study design for bacterial treatment and fecal sample collection. (B) Relative abundance of major phyla between treatment groups across timepoints. (C) Alpha diversity between treatments across time points is represented by the Shannon, Chao1, and Faith’s PD indices. (D) Beta diversity across all timepoints and separated by treatment, represented by Euclidean distances of CLR-transformed data, and weighted unifrac. (E) Differentially abundant taxa (family_genus_sequence#) due to the binary effect of LOX. (F) Differentially abundant taxa due to cumulative doses of LOX.

We next applied PICRUSt2 (71) to infer functional metagenomic profiles from 16S rRNA sequencing data and perform exploratory hypothesis-generating analyses. Principal component analysis of MetaCyc pathway predictions suggests that both probiotic and LOX administration produced global but subtle shifts in predicted metagenomic composition (R^2^= 0.030 and 0.039, respectively), with probiotic administration displaying a dose-response effect, with additional doses increasing effects from mild to moderate strength (R^2^ for cumulative dose = 0.054) (**Fig. 2A**). Although global differential abundance testing did not identify individual pathways significantly altered by LOX administration after statistical modeling, visualization of the top contributing features to both PC1 (37.2% of variance) and PC2 (25.5% of variance) provided insight in the driving forces behind community level differences (**Fig. 2B**). In a targeted exploratory analysis of these top-contributing features to PC1 and PC2, cumulative LOX dosing was uniformly associated with moderate to large effect-size increases (Cohen’s d = 0.60-1.00) by 72 h in pathways associated with fructan degradation, B-vitamin biosynthesis, chorismate metabolism, and lipid synthesis (**Fig. 2C**), with p-values approaching or reaching nominal significance (p=0.04-0.10, with most p<0.06). Notably, the observed effects display a cohesive metabolic trend, as increased pathways include both enzymes for fatty acid synthesis and an essential cofactor (biotin), alongside electron transport chain components (ubiquinone, heme) and respiratory enzymes. This coordinated upregulation of components of biosynthesis suggests a functional shift towards increased anabolic capacity and metabolic activity within the gut microbiota. Additionally, the concurrent increase in fructan degradation pathways further suggests greater metabolic flux toward fermentable substrates supporting SCFA production.

**Fig. 2.**
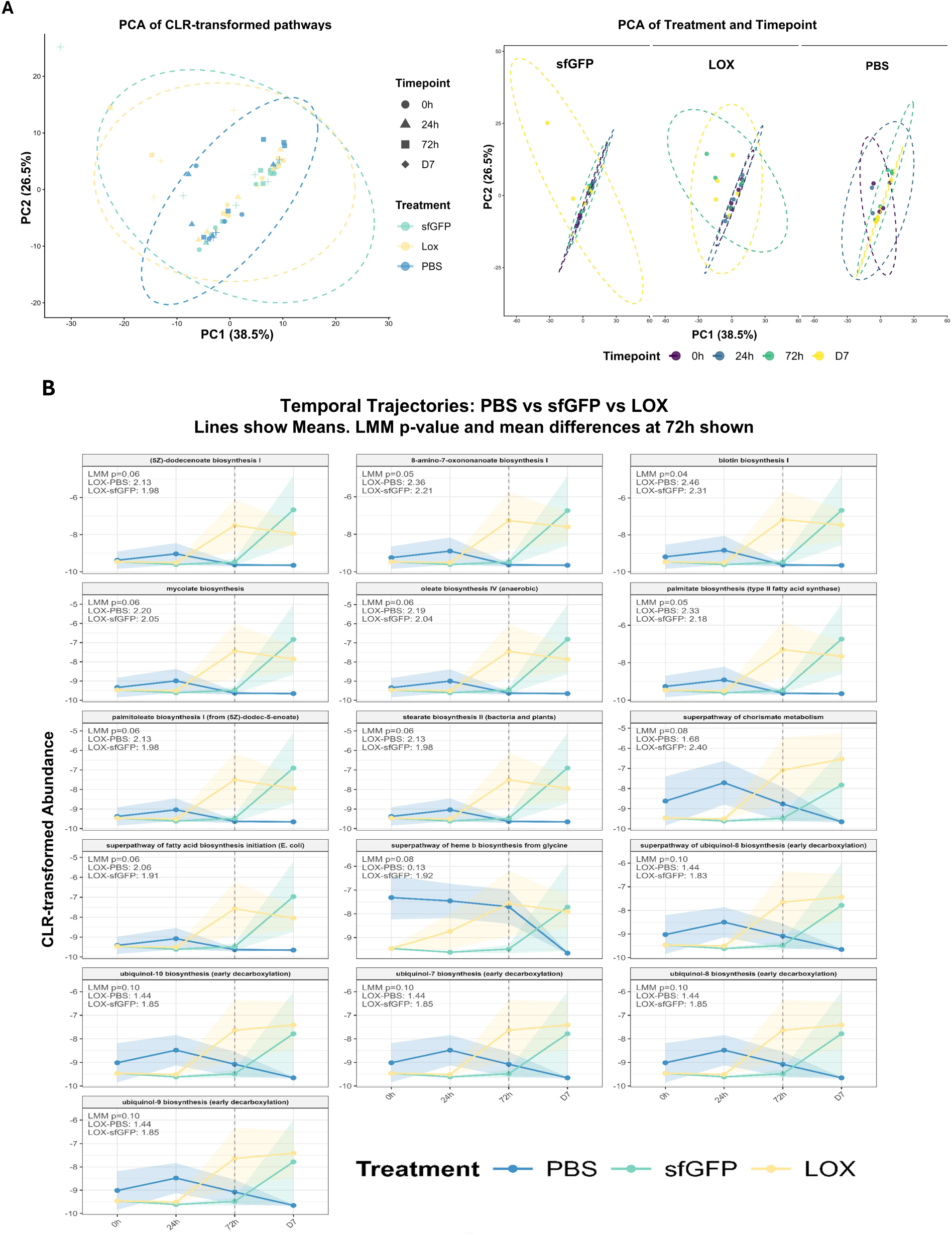
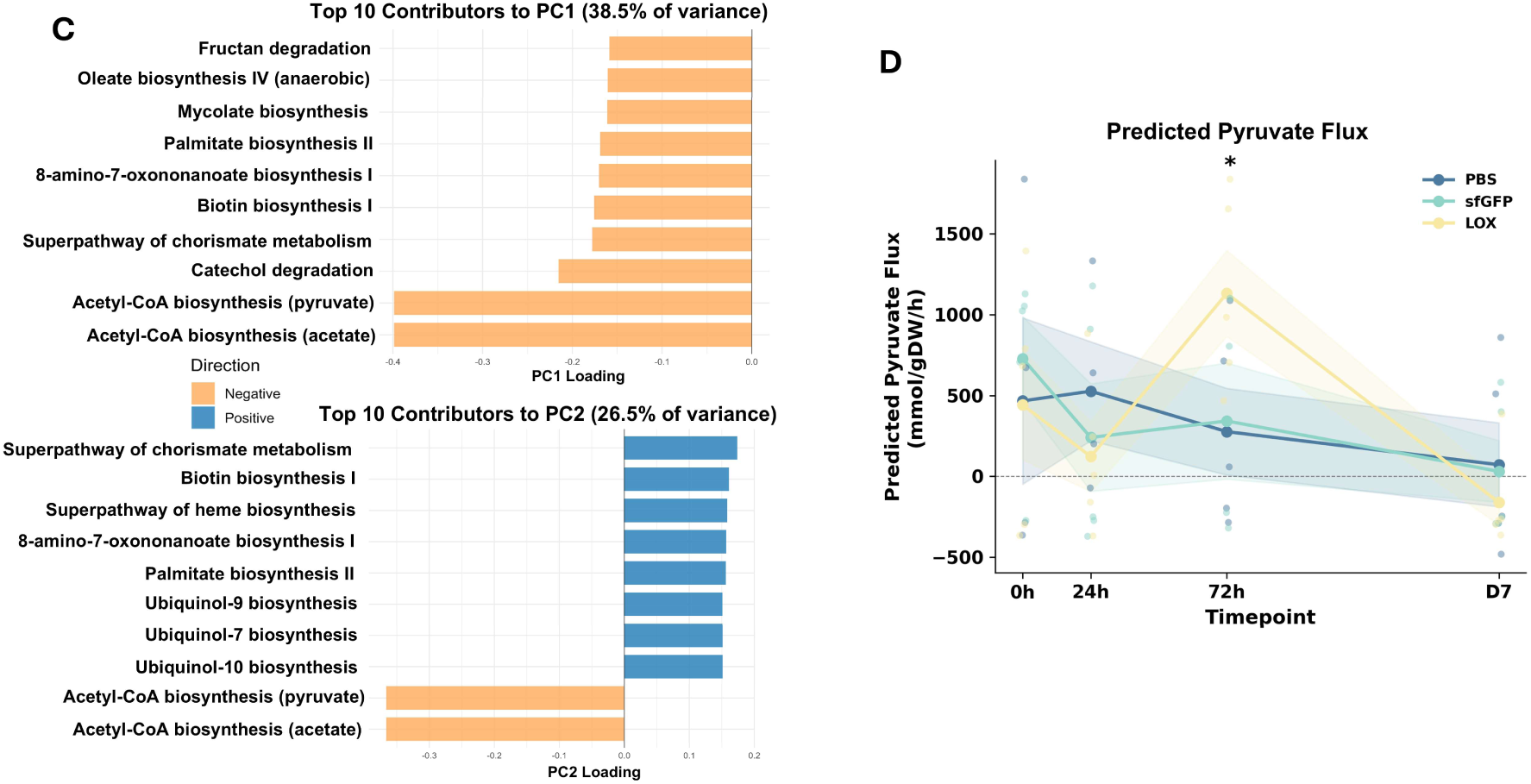
Acute LOX administration drives temporal remodeling of predicted microbiota function. (A) Principal component analysis of CLR-transformed pathway abundances, both combined and separated by treatment. (B) The top 10 contributors to Principal Components 1 and 2, along with their loading values. (C) Temporal trajectories of pathway abundance of 16 candidate pathways (top contributors to PC1 and PC2), which increased due to LOX treatment. (D) MICOM predicted pyruvate flux across timepoints. *indicates significant difference (p<0.05) between LOX and both other groups (sfGFP and PBS).

To assess the functional relevance of this community reorganization, flux-balance analysis with MICOM was used to predict net metabolite flux within the microbial community at steady state. Using genus-level taxonomy from each timepoint, MICOM predicted that the LOX treated community, modeled solely from its observed composition without LOX function, exhibited elevated net pyruvate flux at the 72h timepoint relative to controls (LOX:1131 vs controls: ∼300, p=0.031, **Fig. 2D**), suggesting that LOX treatment reprogrammed the community toward net pyruvate production. Temporal trajectories of other relevant metabolites are plotted in **Fig. S3**. Further, when a GEM for *Bacillus subtilis* with increased lactate uptake (mimicking LOX function) was explicitly included in the model, predicted flux of acetate and formate increased significantly (p=0.0016 and 0.031, respectively for paired within-sample comparison, **Fig. S4,5**), consistent with pyruvate being channeled through pyruvate formate-lyase into acetyl-CoA and formate. These findings support a working hypothesis in which lactate-to-pyruvate conversion by LOX alters carbon availability in the gut, driving coordinated metabolic reprogramming of the resident microbiota toward pyruvate secretion and biosynthetic capacity. Because these inferences are based on functional prediction from 16S data, a modest sample size, and exploratory statistical analyses, they should be viewed as hypothesis-generating and ultimately validated using shotgun metagenomic sequencing and metabolomics in a larger, independent cohort.

### LOX-mediated lactate reduction and microbiome remodeling are temporally associated with transient increases in energy expenditure

Having established that LOX administration, in addition to lowering systemic lactate, is associated with metabolic remodeling of the microbiome at 72 h, we next asked whether these microbial changes coincide with alterations in host energy metabolism. To determine whether the observed calorimetric differences from the treadmill experiment persisted over a physiologically relevant period, we conducted a three-day study in a Comprehensive Lab Animal Monitoring System (CLAMS) using the same pretreatment schedule, with additional doses administered throughout the three-day period (five total doses). When data were analyzed via LME as daily totals, LOX administration increased energy expenditure (β = 1.08, p < 0.05) and glucose oxidation (β = 0.01, p < 0.05), whereas fat oxidation was unchanged. However, analyses at more granular hourly resolution revealed that these observed effects were accentuated on day 1, but diminished on subsequent days for both energy expenditure and glucose oxidation (trt x day 2, trt x day 3 interaction p<0.05), (**Fig. 3A, 3B**).

**Fig. 3.**
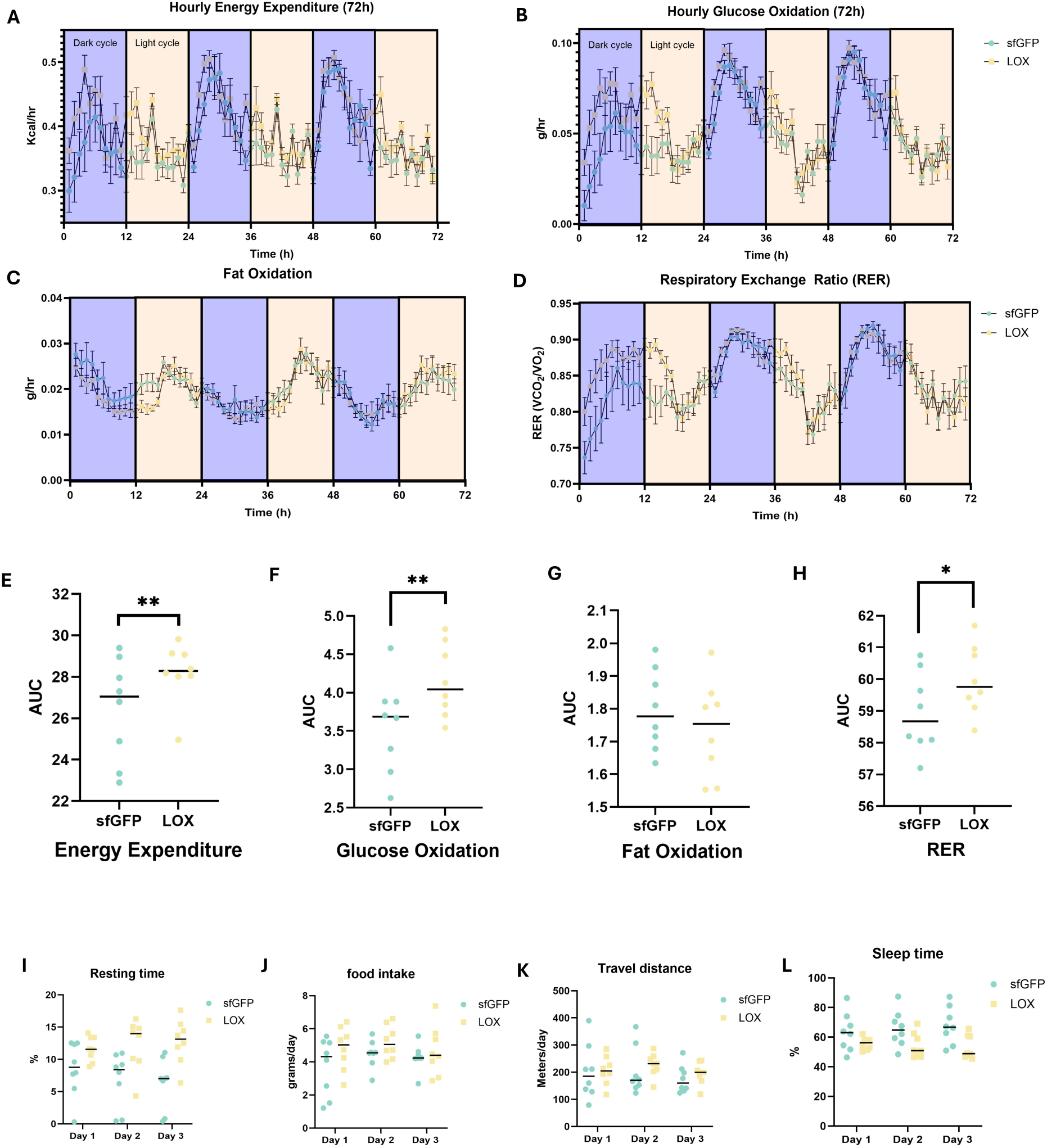
LOX-mediated lactate reduction and microbiome remodeling temporally associate with transient increases in energy expenditure. Hourly values for a 72h period (separated by light/dark cycles) for (A) energy expenditure, (B) glucose oxidation, (C) fat oxidation, and (D) respiratory exchange ratio (RER). Area under the curve for the 72h period for (E) energy expenditure, (F) glucose oxidation, (G) fat oxidation, and (H) RER. Daily total values for (I) resting time, (J) food intake, (K) travel distance, and (L) sleep time.

Behavioral monitoring indicated that LOX-treated mice displayed increased fine movement (β = 7.43, p = 0.046, **Fig. 3**) and resting time (β = 4.96, p = 0.019, **Fig. 3I**), along with reduced sleep time (β = −11.83, p = 0.031) compared to sfGFP. With that said, total travel distance was unchanged (p = 0.617, **Fig. 3K**). Resting time progressively increased over the 3-day period (trt × day3: β = 3.36, p = 0.033, **Fig. 3I**), suggesting progressive behavioral adaptation. Collectively, these findings suggest that effects related to energy expenditure may be attributed to a heightened state of arousal or restlessness during acclimation to a novel environment, as evidenced by increased small movements and reduced rest time, accompanied by decreased sleep. To examine whether the observed metabolic shifts translated to functional improvements, we conducted maximal exercise testing with concurrent indirect calorimetry.

### LOX administration modulates substrate utilization at rest but does not improve endurance performance

Given the strong associations between lactate threshold and peak endurance exercise capacity (49), we tested whether the LOX-induced metagenomic remodeling and/or systemic lactate reductions were capable of improving treadmill running performance in C57BL/6 mice. Our hypothesis was that an increase in biosynthetic activity of the microbiota and a reduction of systemic lactate would lead to increased fuel availability and reduced glycolytic inhibition, resulting in improved endurance performance. Following the same acute treatment schedule as prior experiments, we conducted a test of maximal endurance exercise capacity. Half of the mice in this experiment underwent indirect calorimetry measurements before, during, and after running to exhaustion to characterize substrate utilization and provide further insights into the metabolic alterations caused by LOX administration at rest in a novel environment, during the run to exhaustion, and in the immediate recovery period. Additionally, blood lactate measurements were taken prior to and after treadmill running, but post run sample timing was slightly different between the two treadmills due to the need to collect calorimetry data on the 5-minute recovery period (**Fig. 4A**). Before exercise, LOX treated mice showed significantly reduced blood lactate (**Fig. 4C**), which was associated with reduced glucose oxidation, and showed an insignificant tendency toward elevated fat oxidation (**Fig. 4E**). These effects, however, did not translate to an increase in endurance exercise performance or VO_2_ max (**Fig. 4B**). While LOX treated mice displayed elevated energy expenditure and fat oxidation at rest and during low intensity exercise, both groups responded equivalently to increased exercise intensity (**Fig 4D, 4E**). While all post-exercise time points showed a tendency toward reduced blood lactate in the LOX group, none of these decreases were statistically significant. These data indicate that acute LOX-mediated microbiota remodeling and lactate reduction are sufficient to increase resting energy expenditure and fat oxidation, but do not enhance endurance performance or alter bioenergetics during high-intensity exercise. This dissociation may reflect the redistribution of blood flow away from the gut toward working muscle. Alternatively, the rapid host-driven surge in lactate production during intense exercise likely outpaces the lactate-metabolizing capacity of gut-localized bacteria. These mechanistic hypotheses, and our conclusions, remain tentative given the brief measurement window and modest sample size, and will require validation in longer-term, adequately powered studies.

**Fig. 4.**
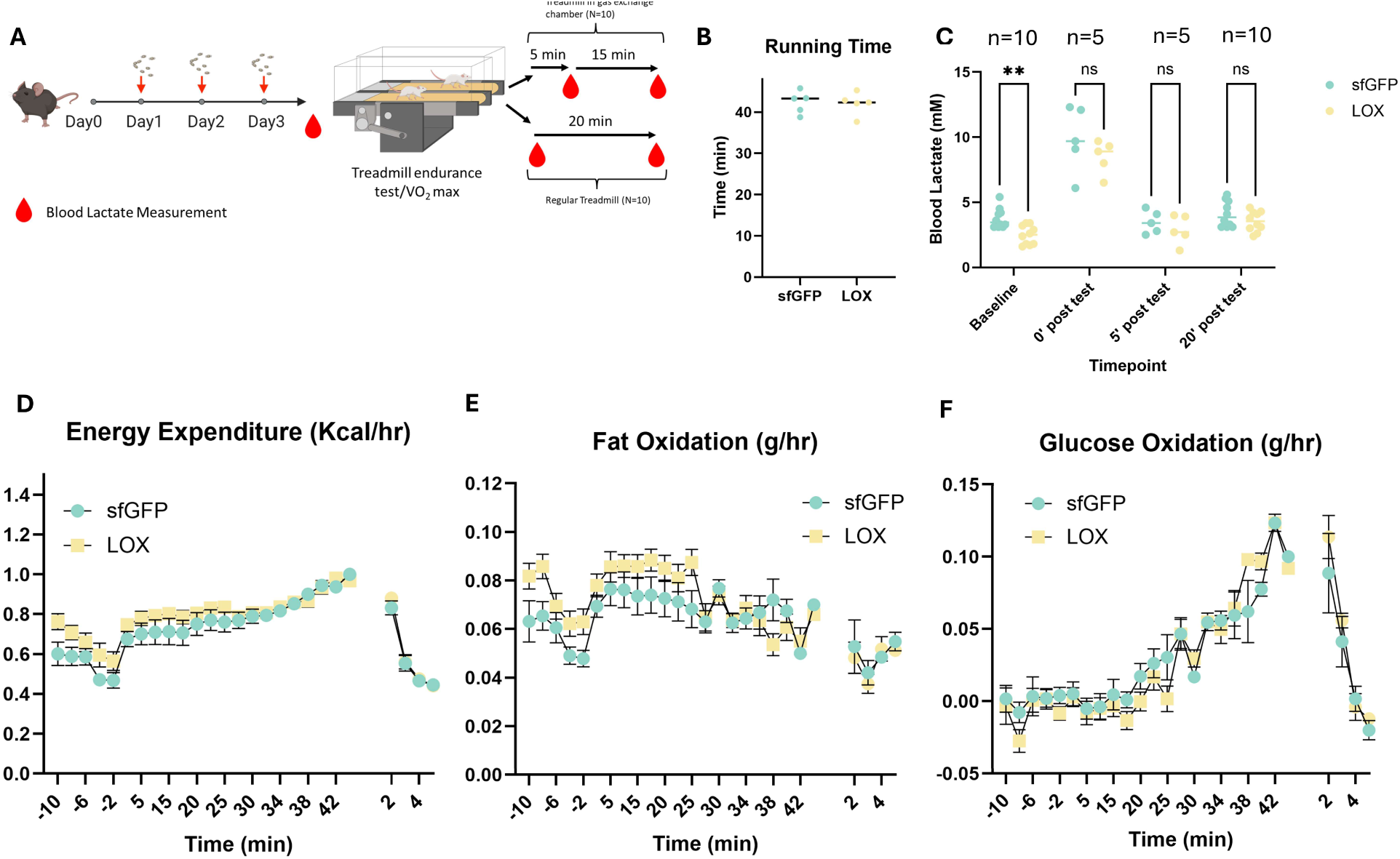
LOX administration modulates substrate utilization at rest but does not improve endurance performance. (A) Schematic diagram of the study design of treadmill experiments. (B) Time to exhaustion in the maximal treadmill endurance capacity test. (C) Blood lactate concentrations at various timepoints throughout the trial periods. Estimated (D) energy expenditure, (E) fat oxidation, and (F) glucose oxidation 10 minutes prior to (negative values), during (first set of positive values), and for 5 minutes after (after gap) the treadmill endurance test.

### Acute LOX administration improves thermoregulation and prolongs survival during LPS-induced sepsis

Having established that acute LOX administration transiently increases host energy expenditure, we next tested whether this metabolic augmentation confers protection during acute inflammatory challenge. To investigate whether the acute increase energy expenditure induced by LOX administration could help the host meet metabolic demands and improve survival rates, we utilized the same acute pretreatment design before induction of LPS-induced sepsis (Fig. 6A). 24h after LPS dosing, LOX treated mice showed marginally reduced blood lactate (p=0.0767) and displayed blunted hypothermia (p=0.029). Body weight loss was equivalent between groups (p=0.74). Given the hypothesis that LOX-mediated metabolic support would primarily influence early survival during the acute inflammatory period, we employed the Gehan-Breslow-Wilcoxon test as the primary analysis, which weights early events more heavily. This test revealed a significant improvement in survival in LOX treated mice over sfGFP (p=0.0479). However, overall survival remained low in both groups, with only 1/22 LOX and 0/21 sfGFP mice surviving to the 5 day endpoint, so the log-rank test for overall survival did not reach significance (p=0.1615).

### Chronic LOX administration reveals homeostatic compensation despite sustained lactate reduction

Given that acute LOX administration resulted in transient metabolic effects without improvement in exercise performance, we next evaluated whether chronic treatment could produce sustained therapeutic benefits in a disease model. To determine whether these observed increases in energy expenditure and glucose oxidation resulted in mitigation of the progression of metabolic disease, we started obese mice on a chronic administration paradigm with additional vehicle control (water) and industry benchmark (semaglutide) treatment groups (**Fig. 5A**). Fecal CFU were assessed to confirm sustained presence of probiotics in the gut between treatments, and it was verified that significant CFU were still detectable at day 3 post dosing (**Fig. S6**). As expected, we saw significant mitigation of weight gain due to semaglutide treatment (p=0.001, **Fig. 5C**). However, despite consistently reduced blood lactate in the LOX group compared to all others (**Fig. 5B**), LOX administration did not impact weight gain (**Fig. 5C**). Further, we conducted metabolic cage experiments on LOX and GFP mice at week 6 and did not find differences in daily energy expenditure, fat oxidation, or glucose oxidation (**Fig. 5E-L**). This finding revealed that the acute changes in energy expenditure and bioenergetic fuel usage were transient alterations that were insufficient to overcome homeostatic regulation in obese mice, displaying the biological limitations of single metabolite interventions.

**Fig. 5.**
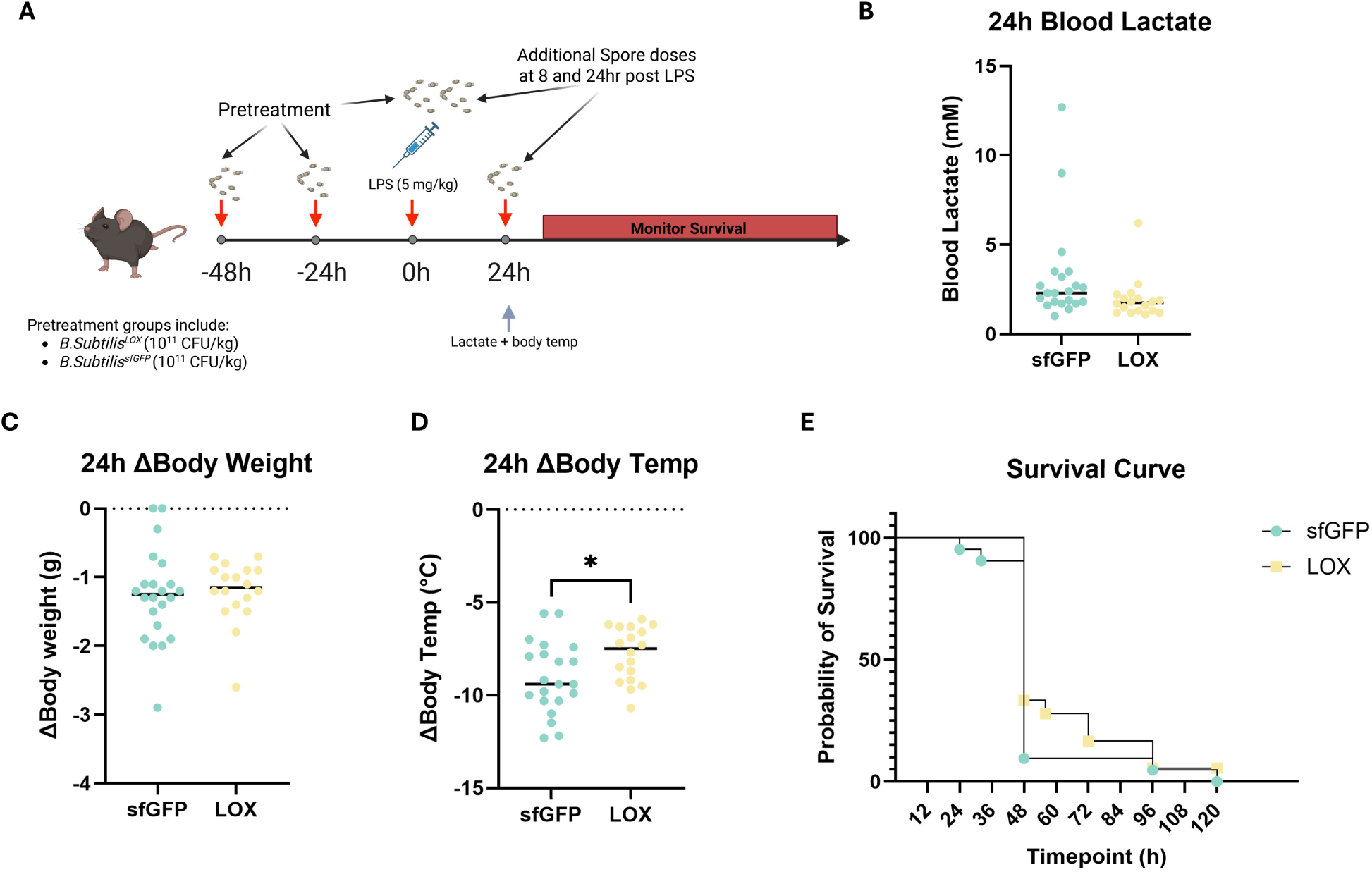
Acute LOX administration improves thermoregulation and survival in endotoxin-induced sepsis. (A) Schematic diagram of study design for sepsis experiments. (B) Blood lactate values 24h post LPS. Changes between the 0h and 24h timepoint in (C) body weight, and (D) body temperature. (E) Survival of each treatment group following LPS treatment.

**Fig. 6.**
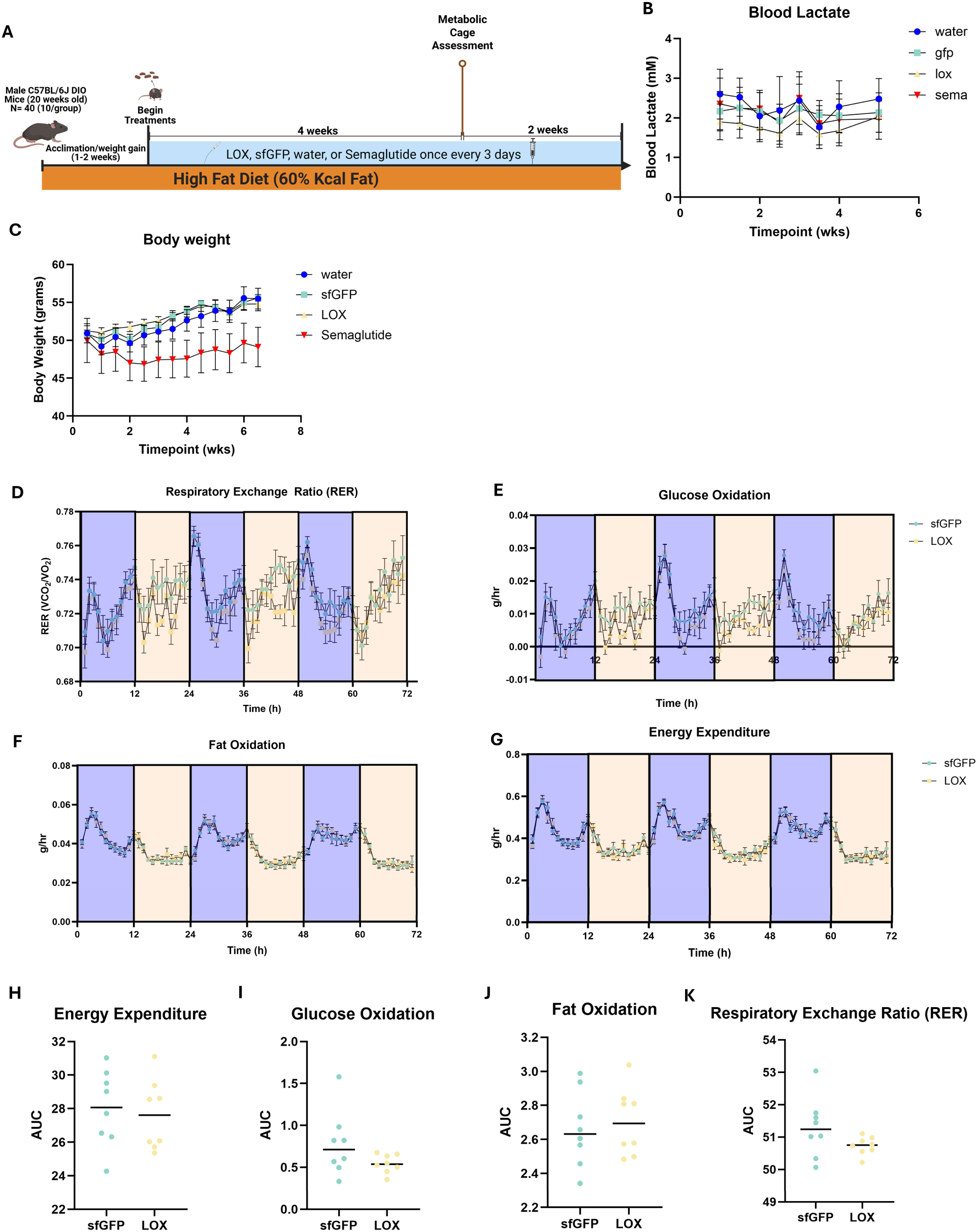
Chronic LOX administration persistently lowers blood lactate but does not sustain metabolic influence or mitigate diet induced obesity. (A) Schematic diagram of study design for obesity experiments. (B) Resting blood lactate values throughout the trial period, (C) Body weight over time. Hourly values for a 72h period (separated by light/dark cycles) for (D) energy expenditure, (E) glucose oxidation, (F) fat oxidation, and (G) respiratory exchange ratio (RER). Area under the curve for the 72h period for (H) energy expenditure, (I) glucose oxidation, (J) fat oxidation, and (K) RER.

## Discussion

Here, we evaluate a novel, functionally driven approach to microbiota engineering, centered on rapid, irreversible conversion of lactate to pyruvate through orally administered *Bacillus subtilis* PY79 expressing lactate oxidase. While our previous work showed this method could reduce systemic lactate at rest and during challenge, here we demonstrate its capability to acutely augment host energy expenditure alongside coordinated microbiota remodeling. Bioinformatic prediction tools revealed that lactate reduction and increased whole-body glucose oxidation were accompanied by a functional shift within the microbiota resulting in increased pyruvate flux and biosynthetic activity. Translationally, these shifts resulted in improved thermoregulation and survival in a mouse model of sepsis. These metabolic effects, however, were limited to an acute timeframe and did not confer benefits in a model of chronic metabolic disease. This contrast between chronic failure and acute success suggests a role for lactate in connecting the microbiome to host energetics, while also identifying physiological limitations of single point metabolic interventions, setting the stage for future, comprehensive research into synbiotic approaches.

Given the complex interplay between the microbiota and the host, the transient increase in energy expenditure due to LOX administration could be explained by two non-exclusive mechanisms. First, the increase in energy expenditure, 72h after the initiation of dosing, is directionally consistent with the observed increases in predicted pyruvate flux and biosynthetic pathways of the gut microbiota at the 72h timepoint. As such, shifting the local lactate:pyruvate ratio and providing a more preferrable carbon source for certain microbes results in an acute increase in biosynthetic output, such as acetate and formate production, which were observed in MICOM predictions including augmented lactate flux, and are known to fuel elevated host energy expenditure (72–74). Second, host factors that can be influenced by systemic lactate:pyruvate ratio may also be potential driving forces (44, 46, 48, 52, 75). Built on the Randle cycle of substrate competition (76), our data suggest that systemic lactate reduction under homeostatic conditions may acutely reduce the inhibitory regulation of glycolysis and is less influential on basal fat oxidation. With that said, when artificially induced by our LOX probiotics, a reduced baseline lactate did not alter peak endurance capacity, although it is considered a weak predictor in human observational studies (49), supporting the notion that it is not causative but rather reflective of an underlying metabolic phenotype. Future studies using gnotobiotic models will enable this engineered probiotic to serve as a valuable tool in investigating the causal role of systemic lactate:pyruvate ratios in metabolic processes.

Lactate clearance is a widely used prognostic marker for sepsis outcomes, and the gut microbiota influences systemic metabolite pools, suggesting that microbiota based interventions targeted at lactate metabolism may prove beneficial in the context of sepsis (3, 4, 77). Our LPS induced sepsis experiment revealed that acute LOX treatment improved thermoregulation and survival, while marginally reducing systemic lactate during the first 24h of sepsis. Notably, the 72h pre-treatment period means the increased energy expenditure and glucose oxidation observed in calorimetry experiments were already established at the time of LPS challenge, effectively serving as metabolic priming. Given that hypothermia is a metabolic tradeoff made during rodent sepsis to preserve vital bioenergetic substrate for the immune system, this augmented energy expenditure may provide support in meeting the metabolic demands of severe immune responses during sepsis (78). While these early results are encouraging, future experiments using an additional vehicle control group (water or PBS gavage) and with greater degree of mechanistic characterization are needed. In particular, an understanding of metabolite differences in the gut and systemic circulation will help improve on this model for this purpose and translate it to the human condition. At present, it is still unknown whether the reduction in lactate, or microbiota functional remodeling are responsible for improvements in thermoregulation. While MICOM experiments predicted an increased pyruvate flux, characterizing shifts in the metabolome could identify whether beneficial metabolites such as pyruvate, succinate, SCFA, or additional microbially-derived metabolites are being released into circulation to support organ function. However, these data at least suggest that engineered and targeted metabolic interventions may be able to provide immune and metabolic support during critical illness.

PERMANOVA analyses revealed relatively subtle but significant changes in both microbiota composition and predicted functional profile due to both probiotic administration and the inclusion of the LOX enzyme. Targeted exploratory analysis of top contributors to principal components revealed a cohesive and coordinated enrichment of several metabolic pathways with moderate to large effect sizes (Cohen’s d = 0.60-1.00), suggesting LOX administration may be a valuable component of combinatorial synbiotic approaches. The LOX group displayed elevated abundance in pathways associated with increased nutrient availability and biosynthetic capacity, like fructan degradation, B vitamin biosynthesis, and chorismate metabolism. Further, these changes temporally aligned with MICOM community simulation findings which revealed that the LOX treated microbiota displayed an increased pyruvate flux, and that inclusion of LOX imitating functions resulted in increased flux of acetate and formate. These predictions, suggesting export of bioavailable carbon sources, may enable LOX probiotics to serve as *in situ* prebiotics, providing cross-feeding substrates which create a suitable functional environment for the engraftment of beneficial taxa in a synbiotic context. Further, the increased export of acetate, a key substrate for butyrate production, may serve as mechanistic basis for establishing a synbiotic approach designed to augment butyrate production.

While the present data do not allow us to parse between effects due to systemic lactate:pyruvate ratios or microbiota activity, the discrepancy between the effects of acute and chronic administration on host metabolic status suggests host or microbial recalibration and return to homeostatic conditions over time. Chronically elevated blood lactate concentrations are concomitant with many forms of metabolic disease (47), and several preclinical studies have shown that chronic exposure to elevated lactate concentrations can induce metabolic defects such as impaired mitochondrial fatty acid uptake (44), adipose inflammation (46), muscular insulin resistance and inhibition of lipolysis (75). With that said, the present study suggests either 1) that chronic lactate elevation is not required for the development of obesity, or 2) the reductions due to LOX administration are too subtle to influence its progression. The discrepancy between acute and chronic effects demonstrates the resilience and strong adaptive capacity of both host and microbiota metabolism. Prior research has shown lasting metabolic benefits in the context of obesity following administration of natural isolates known to be butyrate producers such as *Roseburia hominis* (73), *Faecalibacterium pausnitzii* (74), *Eubacterium rectale*, and *Clostridium butyricum* (72), suggesting that a consortium approach may be required to overcome homeostatic forces in a sustained manner.

To optimize this strategy, future work must address the methodological limitations of the current study. While the addition of MICOM community simulations provides stronger mechanistic predictions than PICRUSt2 findings alone, both approaches remain computational inferences. Accordingly, metabolomics measurements and metagenomic sequencing are needed to define how lactate-to-pyruvate conversion alters the microbiota and luminal/systemic metabolite pools. Future experiments will interrogate the effects of LOX on the local and systemic metabolome, and the extent and nature of functional microbiota remodeling in the context of sepsis. Further, given that the LOX model is a defined, single-node metabolic perturbation with experimentally quantified kinetic parameters, it can serve as a valuable platform for calibration of computational methods by iterating between computation and physical experiments. Additionally, the role of LOX probiotics in establishing an ecological niche for the potentiated engraftment of butyrate-producing bacteria or other live bacterial therapeutic chassis which prefer pyruvate over lactate as a carbon source, such as *EcAZ* or *Nissle 1917* can be investigated to form engineered consortium approaches (79, 80). While *PY79*’s transient gut residence and potent enzymatic activity may be well-suited for acute interventions such as sepsis support, the use of gut native strains such as *EcAZ, NGF-1*, or *Bacteroides thetaiotaomicron* may be beneficial to achieve sustained functional modulation for chronic applications such as metabolic disease treatment. Collectively, this work demonstrates that engineered probiotics with potent, single functions can serve as not only therapeutic candidates, but also as precision tools for dissecting microbiome-host metabolic interactions. These findings establish a framework for mechanistic discovery via engineered probiotics, and rational design of microbial therapeutic interventions.

## Acknowledgments

This work was supported by NIBIB (R21EB030769) and CDMRP PRMRP (W81XWH-21-1-0016), and NH is supported by NIAID (F32AI194752). WBSGFP was kindly provided by Prof. Xin Yan from Nanjing Agricultural University, China. We would like to thank Zhe Wu and the staff at the Michigan Metabolic, Physiological, and Behavioral Phenotyping Core for their generosity in sharing materials and their facilities for the completion of these studies. Phenotyping core equipment and work is supported by MMPC-Live funding (1U2CDK135066). Generative AI was used to aid in the assembly of coding pipelines for bioinformatic analyses, and in ensuring clarity and flow in writing (Claude Sonnet/Opus 4.5/4.6 and Gemini 3). Following the use of these tools, the authors reviewed and edited output as needed and take full responsibility for the accuracy and integrity of the final content and data.

## Author contributions

JL conceived the idea for the LOX model and supervised the experiments. NH designed experiments and statistical modeling, conducted bioinformatic analyses, interpreted outcomes, and directly oversaw the completion of animal experiments and spore preparation, with the help of other lab members and core staff. NY, CF, and MJ assisted with the animal work and spore preparation. NQ provided guidance on design and interpretation of metabolic phenotyping experiments. All authors contributed to the discussion of the results and the writing and editing of the manuscript.

## Data availability statement

The data that support the findings of this study are available from the corresponding author, JL, upon reasonable request, but are also available in online repositories. In vivo data is available via Zenodo (https://doi.org/10.5281/zenodo.18343774), and sequencing database is located in an NCBI SRA database (PRJNA1406803)

