## Supplemental figures for "Engineered Lactate Catabolizing Probiotics Reveal Timescale Dependent Microbiome-Host Metabolic Coupling"

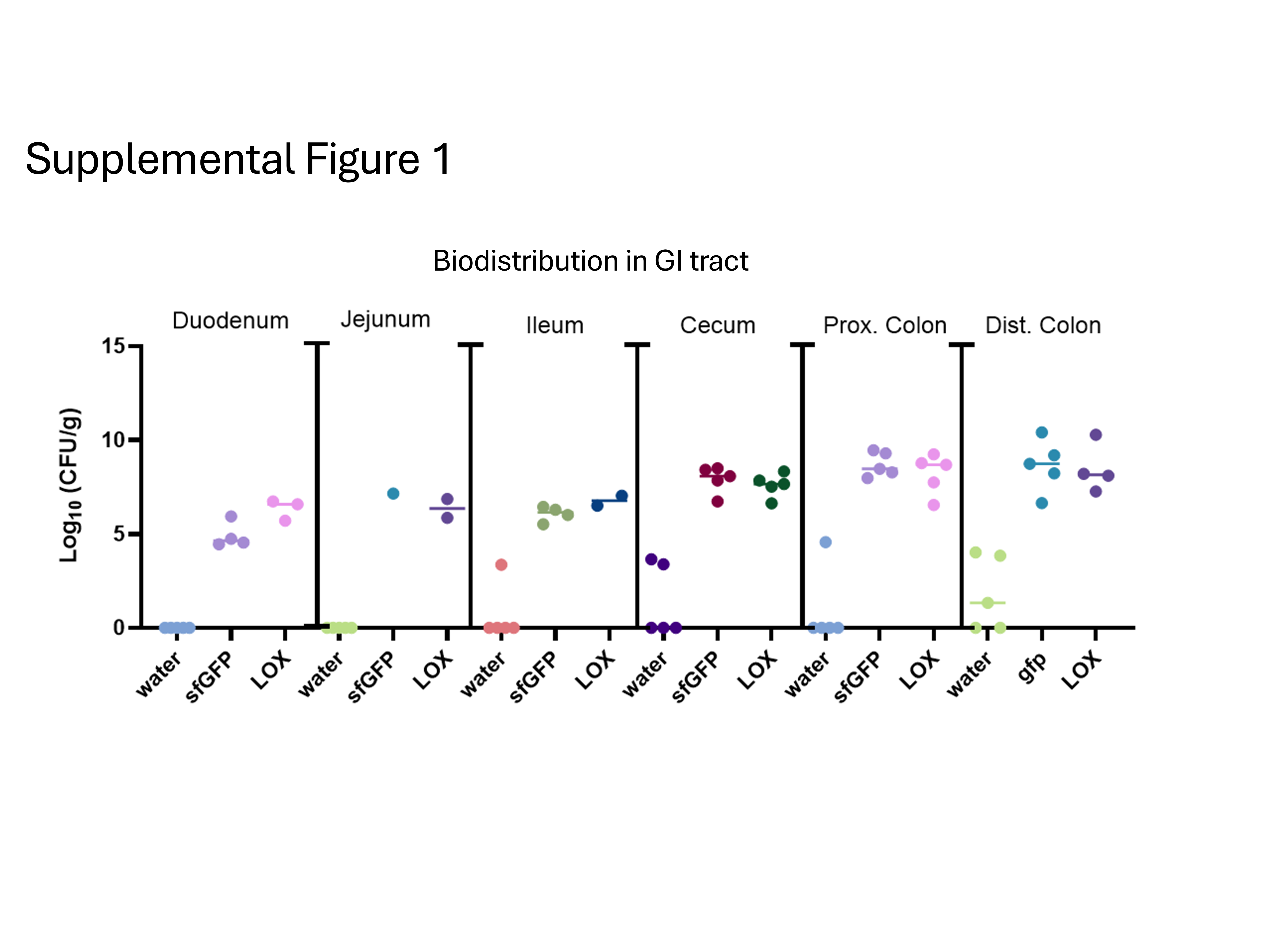
 **Supplemental Figure 1. Biodistribution in the gastrointestinal tract.** Following LOX administration, contents of intestinal regions were removed and plated on LB agar with 5µg/mL chloramphenicol for CFU quantification.

**Supplemental Figure 2. Top 50 significant features in differential abundance analysis.** Differentially abundant ASVs, and the contributing independent variable as identified in differential abundance testing in 16s sequencing data.

**Supplemental Figure 2. Top 50 significant features in differential abundance analysis.** Differentially abundant ASVs, and the contributing independent variable as identified in differential abundance testing in 16s sequencing data.

**Supplemental Figure 2. Top 50 significant features in differential abundance analysis.** Differentially abundant ASVs, and the contributing independent variable as identified in differential abundance testing in 16s sequencing data.

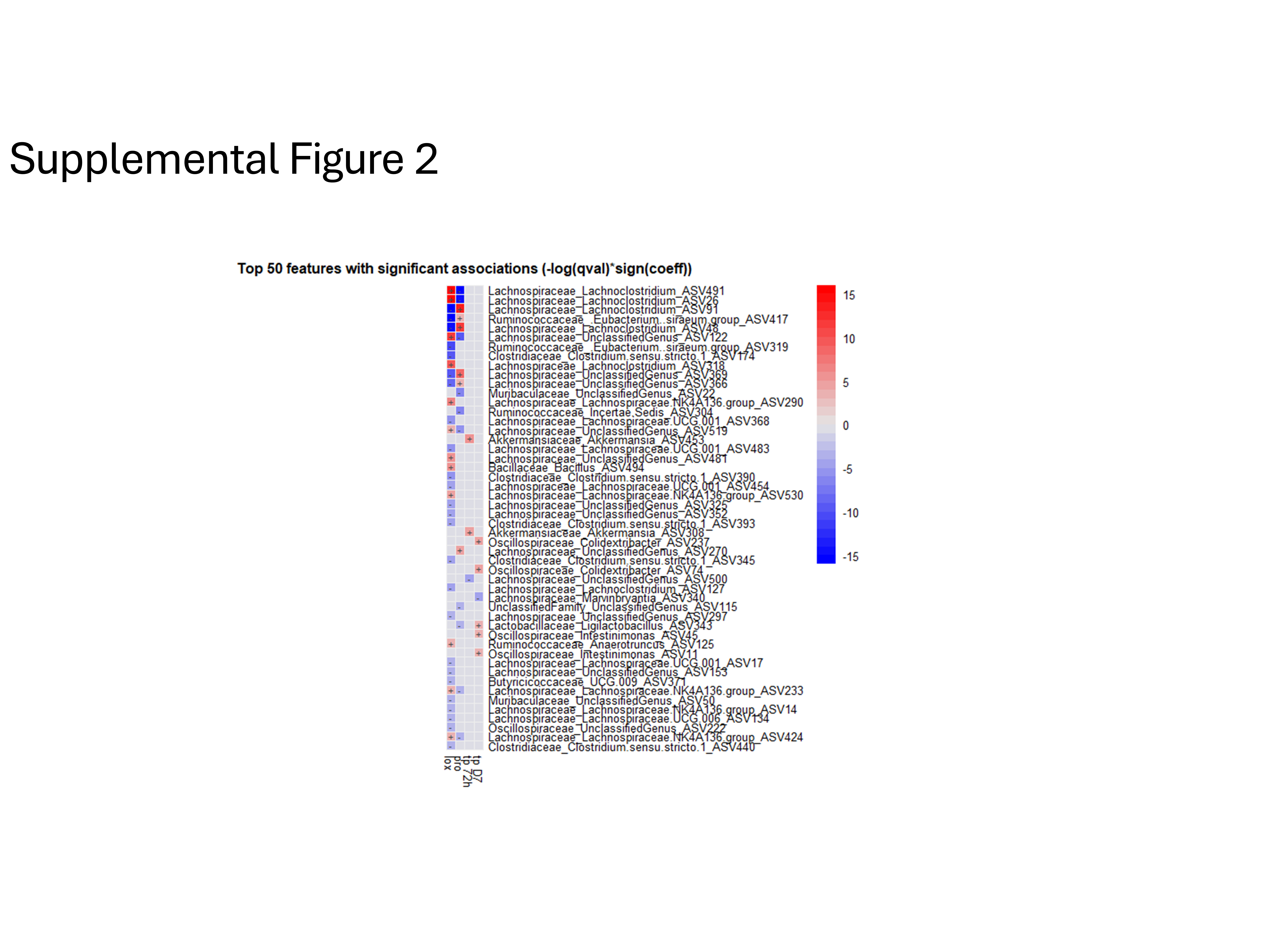
 **Supplemental Figure 2. Top 50 significant features in differential abundance analysis.** Differentially abundant ASVs, and the contributing independent variable as identified in differential abundance testing in 16s sequencing data.

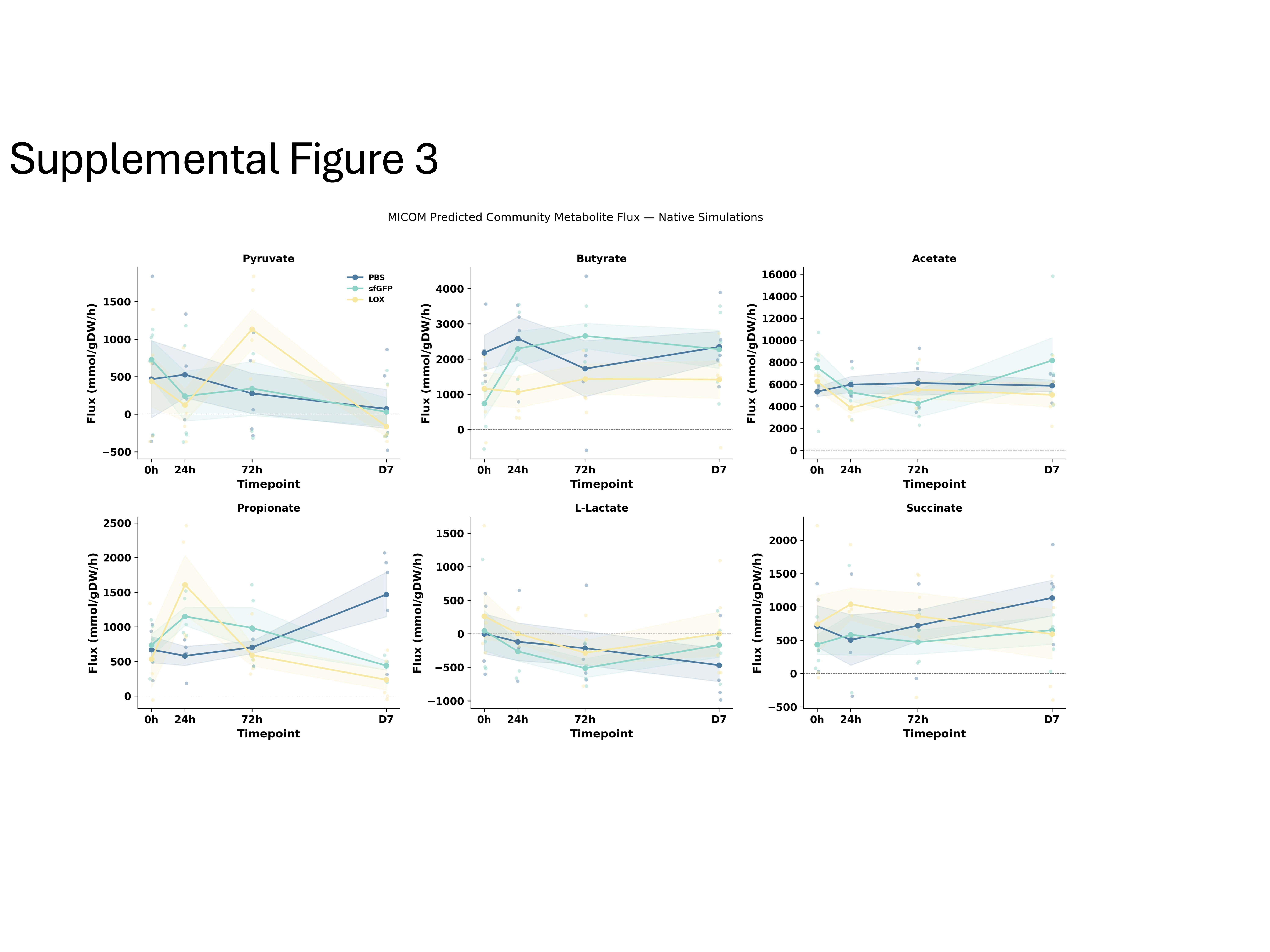
 **Supplemental Figure 3. MICOM-predicted community metabolite flux across timepoints in native simulations.** Predicted net community exchange fluxes for six metabolites (pyruvate, butyrate, acetate, propionate, L-lactate, succinate) across four timepoints (0h, 24h, 72h, Day 7) in native MICOM simulations incorporating the observed microbiome composition for each sample. Lines represent group means; shaded bands indicate ±SEM. Individual sample values are shown as jittered points. PBS, n=20; sfGFP, n=19; LOX, n=20. Positive flux values indicate net community production; negative values indicate net consumption.

**Supplemental Figure 4. Isolated enzymatic effect of LOX on community metabolism.** Paired comparison of MICOM-predicted net community exchange fluxes between chassis-only (sfGFP-expressing *B. subtilis* at 3% relative abundance, no added enzymatic activity) and with-LOX (LOX-expressing *B. subtilis* at 3% relative abundance, forced L-lactate uptake of −12.0 mmol/gDW/h) simulations. Only LOX-group samples at 24h and 72h are shown (n=10 pairs). Lines connect matched samples. Large circles represent mean ± SEM. Significance was assessed by paired t-test. ***, p<0.001; **, p<0.01; *, p<0.05; ns, not significant.

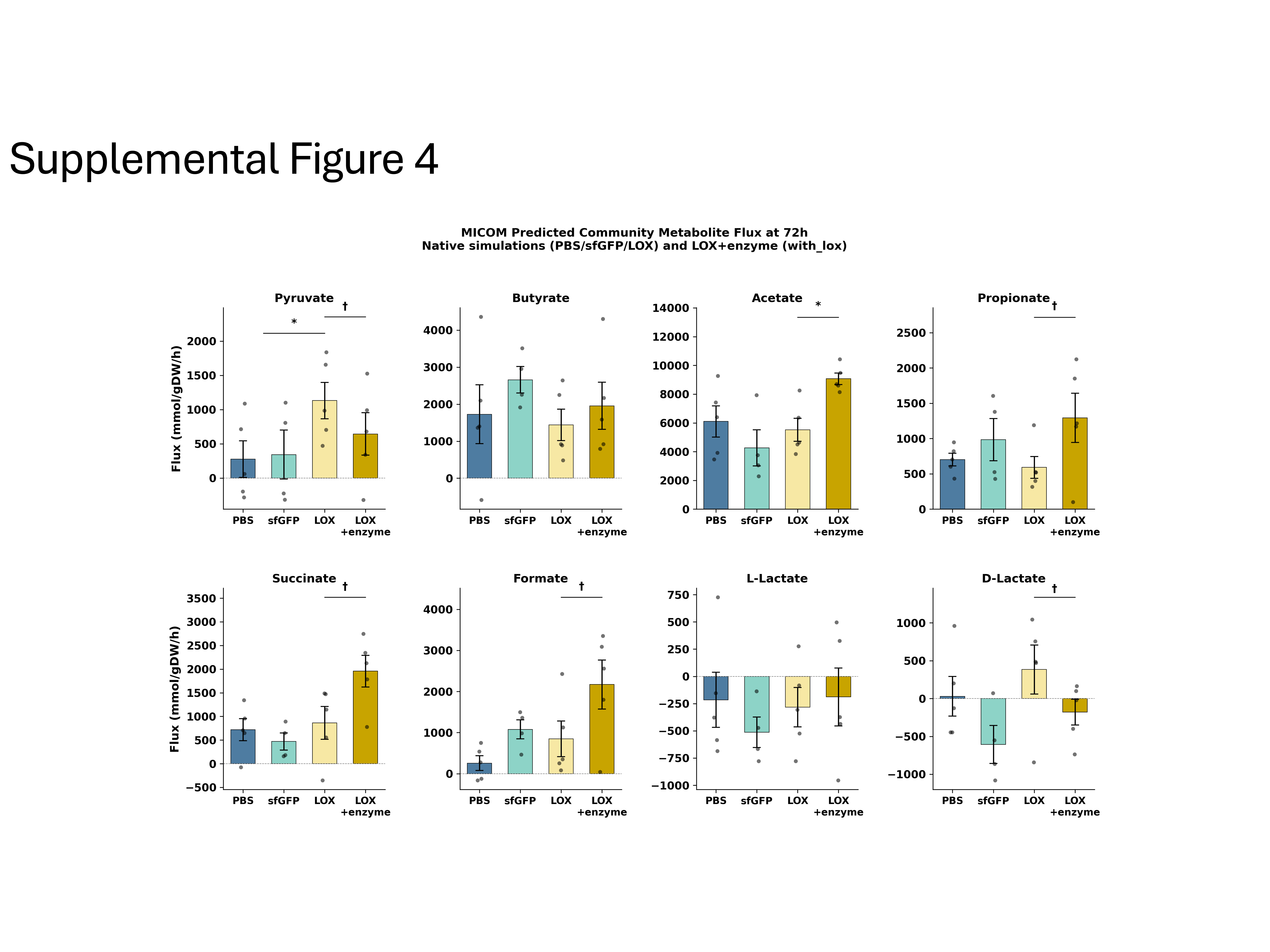
 **Supplemental Figure 4. Isolated enzymatic effect of LOX on community metabolism.** Paired comparison of MICOM-predicted net community exchange fluxes between chassis-only (sfGFP-expressing *B. subtilis* at 3% relative abundance, no added enzymatic activity) and with-LOX (LOX-expressing *B. subtilis* at 3% relative abundance, forced L-lactate uptake of −12.0 mmol/gDW/h) simulations. Only LOX-group samples at 24h and 72h are shown (n=10 pairs). Lines connect matched samples. Large circles represent mean ± SEM. Significance was assessed by paired t-test. ***, p<0.001; **, p<0.01; *, p<0.05; ns, not significant.

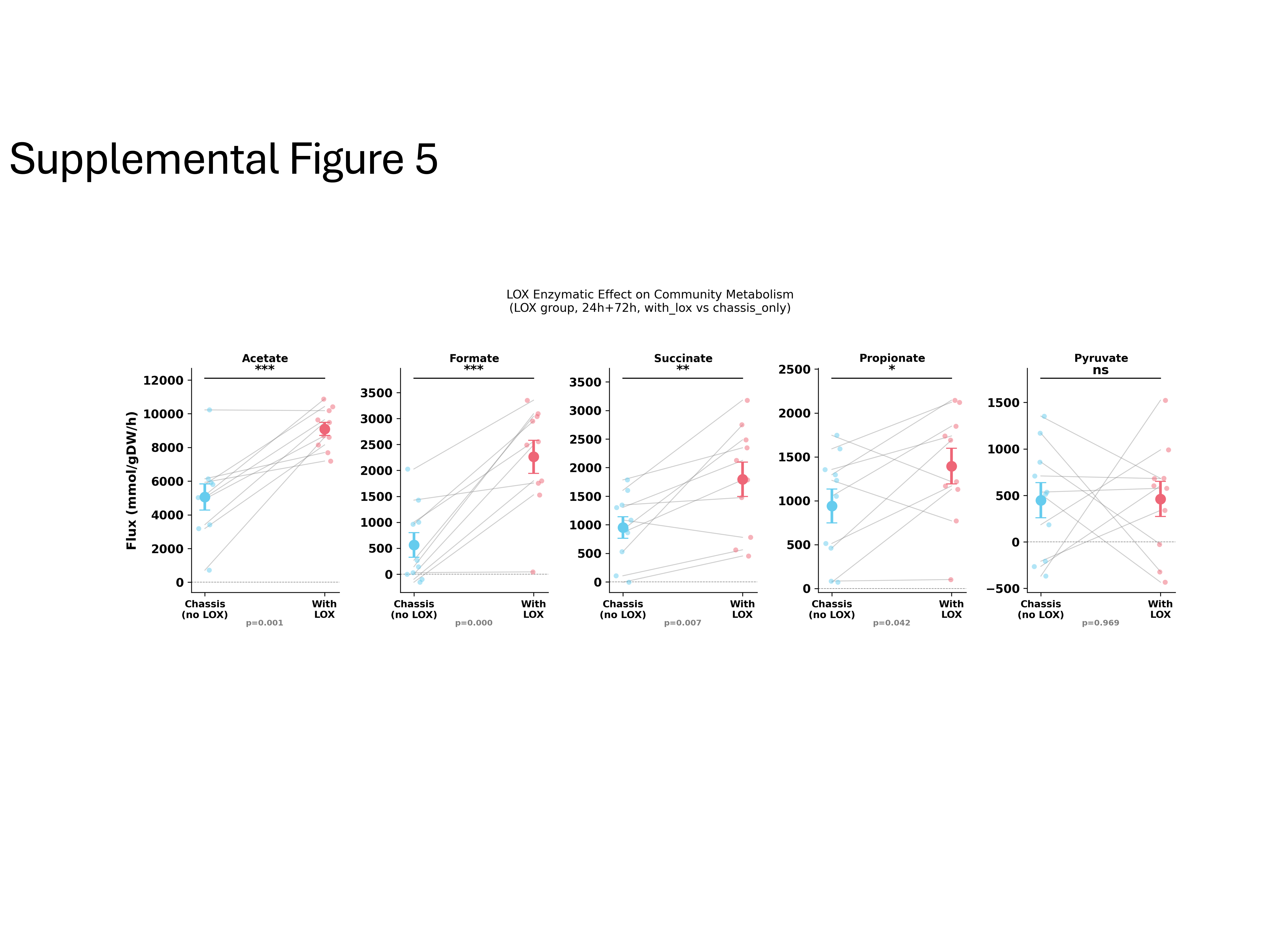
 **Supplementary Figure 5. Cross-sectional MICOM-predicted community metabolite flux at 72h.** Bar plots showing mean ± SEM predicted net community exchange fluxes at the 72h timepoint across treatment groups in native simulations (PBS, sfGFP, LOX) and LOX+enzyme (with-LOX) simulations. Individual sample values are overlaid as scatter points. Significance brackets denote unpaired t-test comparisons between PBS and LOX (native) or between LOX (native) and LOX+enzyme. *, p<0.05; †, p<0.1; no bracket, not significant. LOX+enzyme simulations model the direct enzymatic effect of LOX-mediated L-lactate oxidation on community metabolism and are not directly comparable to the native simulations of the other groups.

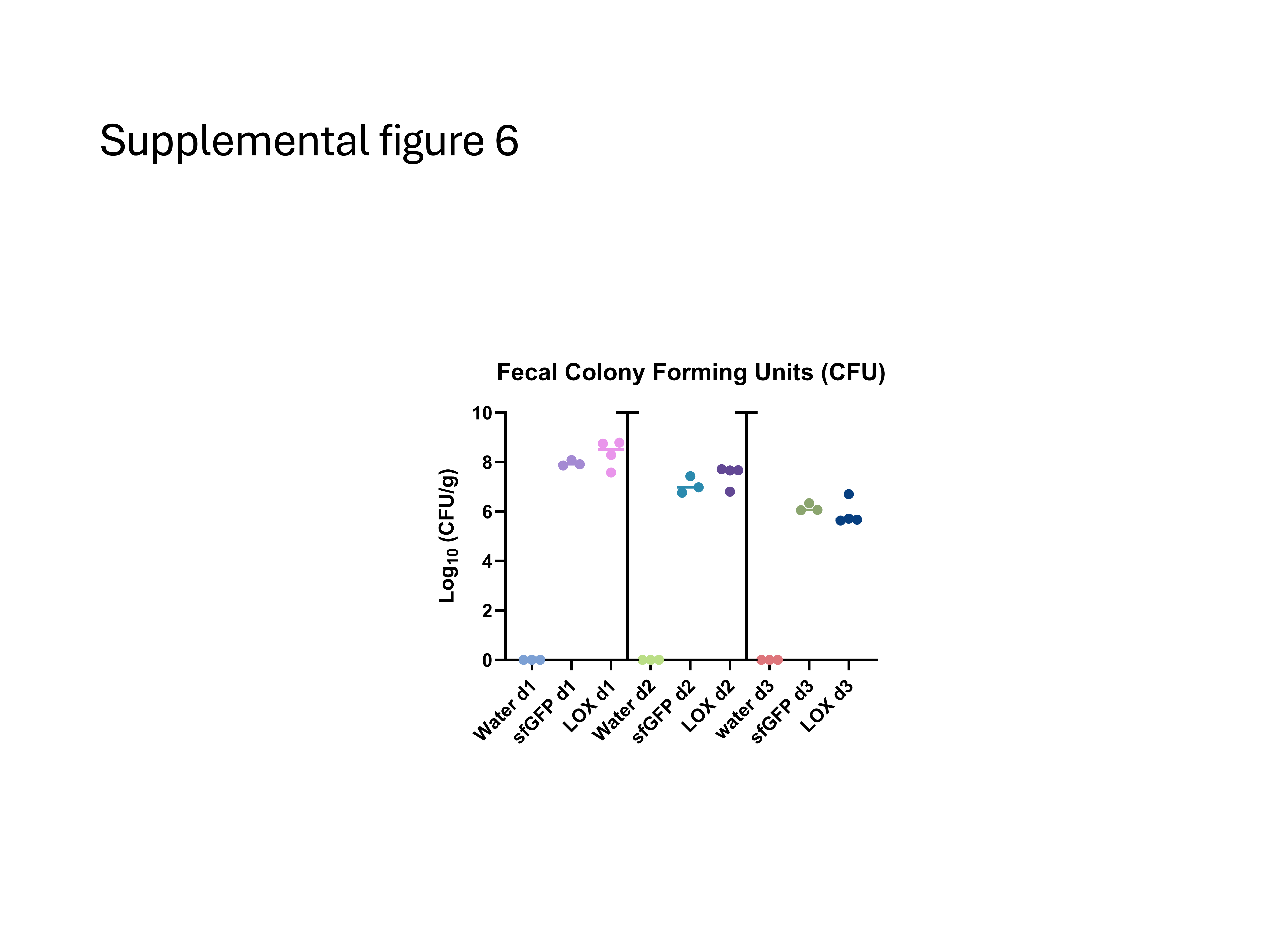
 **Supplemental Figure 6. Temporal kinetics of LOX and sfGFP spores in feces during chronic administration model.** Following an administered dose of respective treatments, fecal samples from LOX, sfGFP, and water treated mice were collected and homogenized in PBS and plated on LB agar with 5µg/mL chloramphenicol for CFU quantification 24h (d1), 48h (d2), and 72h (d3) after administration.
